# Hexosamine biosynthetic pathway integrates circadian and metabolic signals to regulate daily rhythms in protein O-linked N-acetylglucosaminylation

**DOI:** 10.1101/2020.08.18.256636

**Authors:** Xianhui Liu, Ivana Blaženović, Adam J. Contreras, Thu M. Pham, Christine A. Tabuloc, Ying H. Li, Jian Ji, Oliver Fiehn, Joanna C. Chiu

## Abstract

The integration of circadian and metabolic signals is essential for maintaining robust circadian rhythms and ensuring efficient metabolism and energy use. Using *Drosophila* as an animal model, we showed that cellular protein O-linked N-acetylglucosaminylation (O-GlcNAcylation) exhibits robust 24-hour rhythm and is a key post-translational mechanism that regulates circadian physiology. We observed strong correlation between protein O-GlcNAcylation rhythms and clock-controlled feeding-fasting cycles, suggesting that O-GlcNAcylation rhythms are primarily driven by nutrient input. Interestingly, daily O-GlcNAcylation rhythms were severely dampened when we subjected flies to time-restricted feeding (TRF) at unnatural feeding time. This suggests the presence of a clock-regulated buffering mechanism that prevents excessive O-GlcNAcylation at non-optimal times of the day-night cycle. We found that this buffering mechanism is mediated by glutamine-fructose-6-phosphate amidotransferase (GFAT) activity, which is regulated through integration of circadian and metabolic signals. Finally, we generated a mathematical model to describe the key factors that regulate daily O-GlcNAcylation rhythm.

## INTRODUCTION

Circadian clocks exist in organisms from all domains of life. These endogenous timers perceive daily rhythms in environmental and nutrient signals and control the timing of physiological and metabolic processes to optimize energy homeostasis (Panda, 2016; Cox and Takahashi, 2019; Patke et al., 2020). At the behavioral level, feeding-fasting cycles exhibit robust daily rhythms that are regulated by the circadian clock (Xu et al., 2008; Cedernaes et al., 2019). At the molecular level, circadian clocks regulate the rhythmicity of cellular metabolic processes to anticipate feeding-induced nutrient influx to ensure efficient metabolism and energy use (Bass and Takahashi, 2010; Eckel-Magan and Sassone-Corsi, 2013; Perelis et al., 2015; Panda, 2016). The circadian clock has been shown to drive daily rhythmic expression of metabolic genes involved in glycolysis, pentose phosphate pathway, gluconeogenesis, lipid oxidation and storage (Bass and Takahashi, 2010; Panda, 2016; Guan et al., 2020). Once metabolized, macronutrients such as sugars, amino acids and lipids, or the lack of nutrients during fasting period can in turn regulate appropriate nutrient sensing signaling pathways, such as those regulated by insulin/target of rapamycin (TOR), adenosine-monophosphate-activated protein kinase (AMPK), glucagon, adipokines, and autophagy, to orchestrate downstream physiological functions (Eckel-Magan and Sassone-Corsi, 2013; Efeyan et al., 2015; Panda, 2016). The coordinated interplay between cellular processes that are regulated by the circadian clock and those regulated primarily by metabolic signaling is essential for maintaining the robustness of circadian physiology and organismal health (Longo and Panda, 2016; Bass, 2017).

Rhythmic nutrient signals from clock-controlled feeding activity not only feedback to maintain the robustness of the circadian oscillator through metabolic regulation, but can also feedforward to regulate rhythmicity of cellular processes beyond the circadian oscillator (Efeyan et al., 2015; Longo and Panda, 2016). There are many examples illustrating metabolic regulation of key components of the circadian oscillator (Martinek et al., 2001; Travnickova-Bendova et al., 2002; Lamia et al., 2009; Asher et al., 2008; Asher et al., 2010; Nakahata et al., 2008; Dang et al., 2016; Ramanathan, et al., 2018). For example, AMPK promotes the degradation of CRYPTOCHROME 1 (CRY1), a key component of the mammalian clock (Lamia et al., 2009), while in *Drosophila*, GLYCOGEN SYNTHASE KINASE 3β (GSK3β) facilitates nuclear entry of the PERIOD-TIMELESS (PER-TIM) transcriptional repressor complex to regulate circadian period length and output (Martinek et al., 2001). Beyond the circadian oscillator, daily oscillation of reactive metabolites can impose timely regulation on transcriptional and chromatin dynamics. Metabolism modulates the global chromatin landscape by rhythmically providing the essential substrates, such as acetyl-CoA for acetylation and S-adenosyl methionine for methylation (Etchegaray et al., 2003; Ripperger and Schibler, 2006; Xia et al., 2015; Etchegaray and Mostoslavsky, 2016; Mauvoisin et al., 2017). Furthermore, metabolites can serve as cofactors to directly affect activities of chromatin remodelers and histone modifiers to regulate the circadian transcriptome. Sirtuins (SIRT), a class of proteins with NAD^+^-activated deacetylase activities, have been identified as key integrators of circadian and metabolic signaling due to their roles in regulating key circadian clock proteins and the chromatin landscape (Asher et al., 2008; Nakahata et al., 2008; Sassone-Corsi, 2012; Masri et al., 2014).

Despite significant progress in understanding the coordination between the circadian clock and metabolism in regulating circadian physiology, our knowledge is far from complete. Specifically, emerging data showing that protein O-linked N-Acetylglucosaminylation (O-GlcNAcylation), a nutrient-sensitive post-translational modification (PTM), is highly prevalent and can regulate diverse cellular processes (Hart et al., 2011; Mishra et al., 2011; Yang and Qian, 2017; Hart, 2019) prompted us to hypothesize that rhythmic nutrient influx through clock-controlled feeding-fasting cycles may regulate time-of-day-specific O-GlcNAcylation and cellular protein functions. Uridine diphosphate-N-acetyglucosamine (UDP-GlcNAc) is the end-product of the hexosamine biosynthetic pathway (HBP) and the donor substrate that enables modifications at serine and threonine residues with O-GlcNAcylation. HBP is considered the most generalized sensor for metabolic status as it integrates metabolites from breakdown of glucose, amino acids, lipids, and nucleic acids (Hart, 2019). Together with the fact that there is extensive interplay between O-GlcNAcylation and phosphorylation to regulate cellular protein functions (Hart et al., 2011; Mishra et al., 2011) and recent findings that a significant portion of the phosphoproteomes in mice and flies exhibit daily oscillations (Robles et al., 2017; Wang et al., 2017; Wang et al., 2020), we sought to investigate if protein O-GlcNAcylation represents an important post-translational mechanism that integrates circadian and metabolic regulation to coordinate circadian physiology.

In previous studies, we and others have shown that a number of key clock transcription factors in flies and mice exhibit daily O-GlcNAcylation rhythms that regulate their subcellular localization, activity, and stability (Kim et al., 2012; Kaasik et al. 2013; Li et al., 2013; Li et al., 2019). However, it is not clear whether cellular proteins beyond the circadian oscillator also exhibit daily O-GlcNAcylation rhythms. In this study, we observed that the O-GlcNAcylation of total nuclear proteins oscillates over a 24-hr period in wild type (WT) *Drosophila*. Nuclear protein O-GlcNAcylation rhythms showed strong correlation with rhythms in food intake and HBP metabolites, suggesting that they could be driving protein O-GlcNAcylation rhythms. We then manipulated timing of nutrient input using time-restricted feeding (TRF) to establish causal relationships between feeding time, HBP metabolites, and protein O-GlcNAcylation rhythm. Despite the importance of feeding time, we observed that the phase of protein O-GlcNAcylation rhythms did not simply shift when feeding occurs at unnatural time window. Rather, the amplitude of the daily O-GlcNAcylation rhythm dampens significantly. This suggests that clock-controlled buffering mechanism(s) exist to limit excessive O-GlcNAcylation if animals feed during non-optimal time of the day-night cycle. By characterizing the daily activity of HBP, we found that glutamine--fructose-6-phosphate amidotransferase (GFAT), the rate limiting enzyme of HBP, is regulated by multiple clock-controlled mechanisms to influence HBP output and protein O-GlcNAcylation rhythm. We observed that *gfat2* mRNA is induced by clock-controlled food intake and GFAT enzyme activity is regulated at the post-transcriptional level, likely by time-of-day-specific phosphorylation. In summary, our results provide insights into the role of HBP in integrating circadian and metabolic signals to regulate daily rhythms in cellular protein O-GlcNAcylation to orchestrate circadian physiology. Our results shed light on the health benefits of TRF and deleterious effects of non-optimal meal times, which are common in modern society.

## RESULTS

### O-GlcNAcylation of nuclear proteins exhibits daily rhythmicity

A number of key circadian clock proteins have previously been observed to exhibit daily rhythms in O-GlcNAcylation that modulate their time-of-day-specific functions (Kim et al., 2012; Kaasik et al., 2013; Li et al., 2013; Li et al., 2019). We hypothesized that this phenomenon may be more widespread and could serve as an important mechanism that regulates daily rhythms in protein structure and function. Using chemoenzymatic labeling (Khidekel et al., 2003), a strategy we have previously employed to examine PERIOD (PER) O-GlcNAcylation (Li et al., 2019), we conjugated biotin tags to O-GlcNAc groups on nuclear proteins extracted from wild type (WT; *w^1118^*) male fly bodies. We observed that a collection of proteins, ranging from 37 to 250 kD, are O-GlcNAcylated in the nuclei extracted from body tissues (Fig. 1A, top panel). The band detected in unlabeled samples (Fig. 1A, middle panel) was deemed to be non-specific and excluded during quantification (Fig. 1A and B). Our results showed that O-GlcNAcylation of nuclear proteins exhibited a robust daily rhythm, suggesting that timely metabolic input may regulate the function of cellular proteins through this post-translational modification. O-GlcNAcylation analyses were conducted only on body tissues as we expected them to be more metabolically sensitive (Xu et al., 2008; Xu et al., 2011; Rhoades et al., 2018). In addition, since the labeling method we used cannot discriminate between GlcNAc groups on N-linked and O-linked glycoproteins, only nuclear proteins were analyzed to avoid contamination from N-glycans in the endoplasmic reticulum, the Golgi apparatus and on the cell surface (Spiro, 2002).

**Figure 1.**
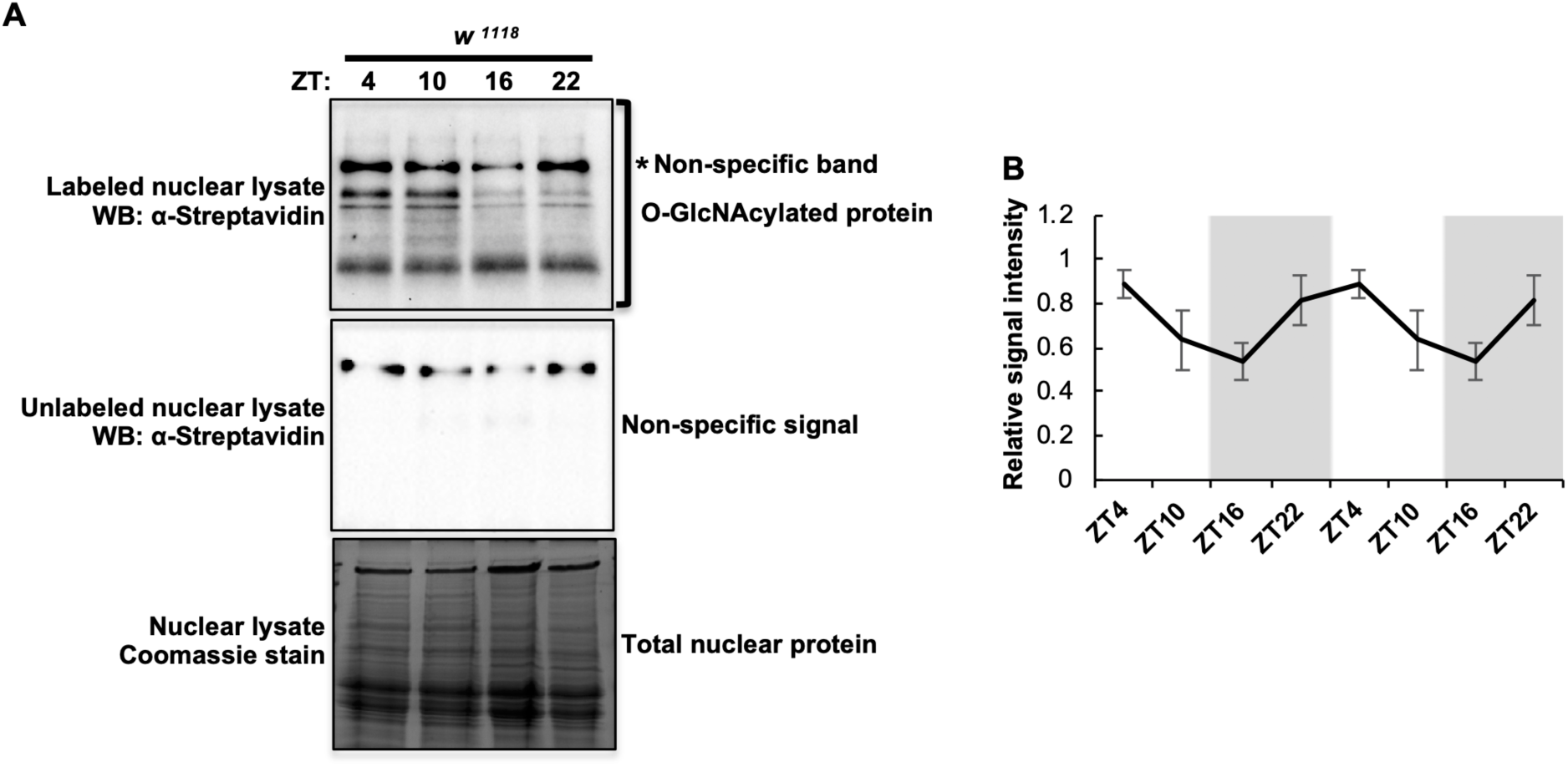
Protein O-GlcNAcylation oscillates over a 24-hour day-night cycle under *ad libtum* condition. (A) Western blots showing daily rhythms in O-GlcNAcylation of nuclear proteins extracted from body tissues of wild type (*w^1118^*) flies fed *ad libtum.* O-GlcNAcylation is detected using chemoenzymatic labeling in combination with Western blotting using α-streptavidin. Unlabeled samples (middle panel) were processed in parallel to labeled samples (top panel) to identify non-specific signal. Total nuclear proteins were stained using Coomassie blue (bottom panel) and were used for normalization. The asterisk (top panel) denotes non-specific signal. (B) Quantification of nuclear protein O-GlcNAcylation in (A). Data are double plotted and presented as mean ± SEM (n=4; p=0.045; RAIN). α-streptavidin signal of the whole lane was quantified, and the non-specific signal was used for background deduction prior to normalization. The grey shading indicates the dark phase of each day-night cycle. ZT: Zeitgeber time (hr).

### Daily rhythmicity of HBP metabolites correlates with feeding-fasting rhythm

Next we investigated whether feeding-fasting rhythm drives daily oscillation of nutrient input and impact the timing of cellular protein O-GlcNAcylation. We first measured timing of daily feeding activities of WT flies using the CApillary FEeder (CAFE) assay (Ja et al., 2007; Xu et al., 2008). Male and female flies were provided with food *ad libitum* and monitored separately over two day-night cycles. Although both sexes exhibit robust feeding-fasting rhythm, peak feeding activity was observed around ZT24 in males as previously reported (Xu et al., 2008) (Fig. 2A) while females exhibited a phase delay relative to males with a peak around ZT12 (Fig. S1A). Because of the observed sexual dimorphism in the timing of feeding activity, only male flies were used in subsequent experiments.

**Figure 2.**
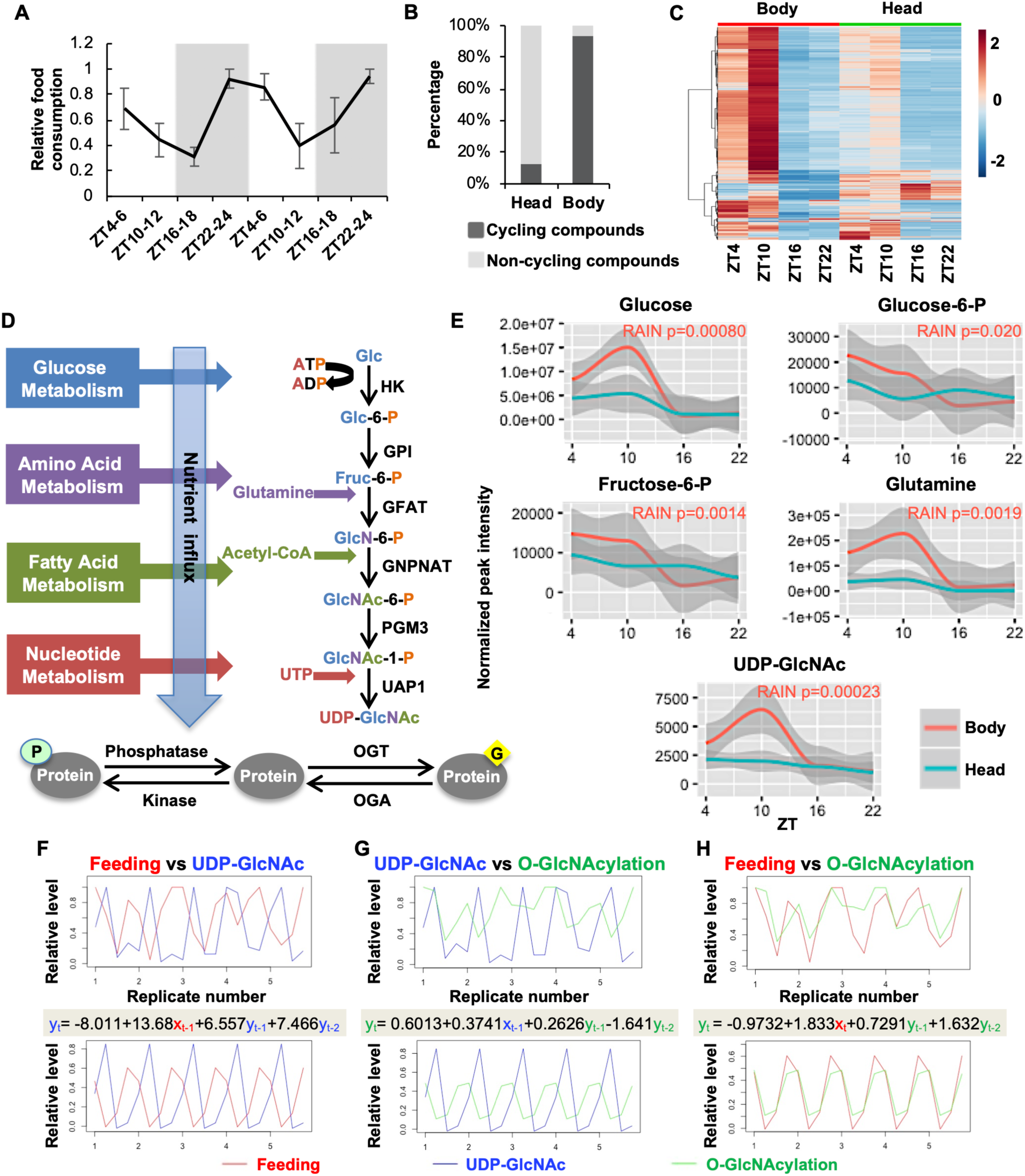
Daily feeding-fasting cycle generates oscillation in metabolites in flies. (A) Feeding rhythm of wild type (*w^1118^*) male flies over two consecutive days as measured by CAFE assay. Data were normalized (peak feeding = 1) and presented as mean ± SEM (n=3, 10 flies per biological replicate; p=0.0014; RAIN). The grey shading indicates the dark phase of each day-night cycle. (B) Percentage of cycling and non-cycling metabolites in male fly heads and bodies detected by GC-MS (n=5 or 6; p<0.05; RAIN). (C) Heat map of all metabolites detected in male heads and bodies (n=5 or 6). Normalized peak intensity of all metabolites were pareto scaled. Each line represents one detected compound. (D) Schematic showing hexosamine biosynthetic pathway (HBP) integrating cellular metabolic status to provide UDP-GlcNAc as output (format of diagram modified from Hart et al., 2011). HK: Hexokinase; GPI: Phosphoglucose isomerase; GFAT: Glutamine--fructose-6-phosphate aminotransferase; GNPNAT: Glucosamine-phosphate N-acetyltransferase; PGM3: Phosphoacetylglucosamine mutase; UAP1: UDP-N-Acetyl glucosamine pyrophosphorylase 1; OGT: O-GlcNAc transferase; OGA: O-GlcNAcase. Glc: glucose; Glc-6-P: Glucose-6-phosphate; Fruc-6-P: Fructose-6-phosphate; GlcN-6-P: Glucosamine-6-phosphate; GlcNAc-6-P: N-acetylglucosamine-6-phosphate; GlcNAc-1-P: N-acetylglucosamine-1-phosphate; UTP: Uridine triphosphate; UDP-GlcNAc: Uridine diphosphate N-acetylglucosamine. (E) Line graphs showing daily oscillations of HBP metabolites in fly bodies and heads (n=5 or 6). Shaded regions represent 95% confidence interval. P values indicating rhythmicity (RAIN) of each of the five metabolites in fly bodies are presented on the upper right corner of each panel. p>0.25 (RAIN) for all HBP metabolites in head samples. (F-H) Cross correlation analysis between rhythms: (F) feeding and UDP-GlcNAc, (G) UDP-GlcNAc and O-GlcNAcylation, (H) feeding and O-GlcNAcylation. Top panels show the observed raw data with all biological replicates, while bottom panels indicate the rhythmic pattern extracted from the raw data using R. The correlation equations are shown between panels with the color of terms corresponding to each curve (For all the equations, R^2^=1).

We harvested fly tissues over a 24-hour period at 6-hour intervals and screened primary metabolites by GC-TOF MS on head and body tissues separately to determine if there is correlation between daily rhythms in feeding and metabolites, especially those in the HBP. We detected 573 metabolites in total, with 167 known and 406 unknown compounds (Dataset S1). Replicates that exhibited 10-fold differences of total peak intensity in comparison to other replicates were excluded in the analysis (Fig. S1B-C). 93.0% of the total compounds cycle in fly bodies while only 12.7% cycle in heads (RAIN p<0.05, Dataset S2) (Fig. 2B). This difference in rhythmicity between head vs body metabolites is likely due to the blood-brain barrier. Similar to the previous circadian metabolomic study conducted in flies (Rhoades et al., 2018), most of the cycling metabolites peak around ZT10, which is subsequent to peak time of food intake, corroborating an anticipated relationship between feeding and nutrient influx (Fig. 2C, Dataset S1). The majority of known compounds, including carbohydrates, amino acids and a small number of lipids, belong to primary metabolic pathways. As we are especially interested in protein O-GlcNAcylation, we targeted the HBP (Fig. 2D) for more in-depth data analysis. HBP is a branch of glycolysis that produces UDP-GlcNAc, the key donor substrate for O-GlcNAcylation. We observed that the first four HBP metabolites and the end product, UDP-GlcNAc, displayed robust 24-hr rhythmicity in body tissues (RAIN p<0.05), while rhythmicity was weak or absent in heads (Fig. 2E, RAIN p>0.25). The oscillation of UDP-GlcNAc, similar to the majority of the detected compounds, is strongly correlated to fly feeding rhythm, with maximum level occurring soon after peak food intake and the trough during the fasting period (Fig. 2F, S2A, B, D and G). Furthermore, we observed strong correlations between the daily oscillations of UDP-GlcNAc vs. O-GlcNAcylation of nuclear proteins and between the timing of feeding activity vs. O-GlcNAcylation rhythm (Fig. 2G-H, S2A-C, E-F and H-I). This supports our hypothesis that feeding-fasting cycle drives the oscillation of UDP-GlcNAc, which in turn promotes rhythmic protein O-GlcNAcylation.

### The circadian clock regulates daily oscillation of protein O-GlcNAcylation via multiple mechanisms

To establish causality between feeding-fasting cycles and rhythmic protein O-GlcNAcylation, we manipulated timing of food intake by time-restricted feeding (TRF) and monitored changes in O-GlcNAcylation patterns in fly body tissues. We divided WT flies into three feeding groups (Fig. 3A): (1) The *ad libitum* (AL) group had food available at all times; (2) the natural feeding (RF21-3) group was only fed between ZT 21 to ZT3, the natural fly feeding time according to our CAFE assay (Fig. 2A); (3) the mistimed feeding group (RF9-15) was fed 12 hours out of phase (Xu et al., 2011). We expect that with a limited feeding window in the TRF groups, the amplitude of the O-GlcNAcylation rhythm may be higher in comparison to that in the AL group in which feeding is relatively less consolidated. We first used the CAFE assay to confirm that the feeding groups consumed similar amount of food daily (Fig. 3B). We then compared the feeding amount in TRF groups to that of the AL group during their respective feeding window to show that the TRF treatments indeed consolidated their daily food intake into the 6-hour feeding window (Fig. 3C).

**Figure 3.**
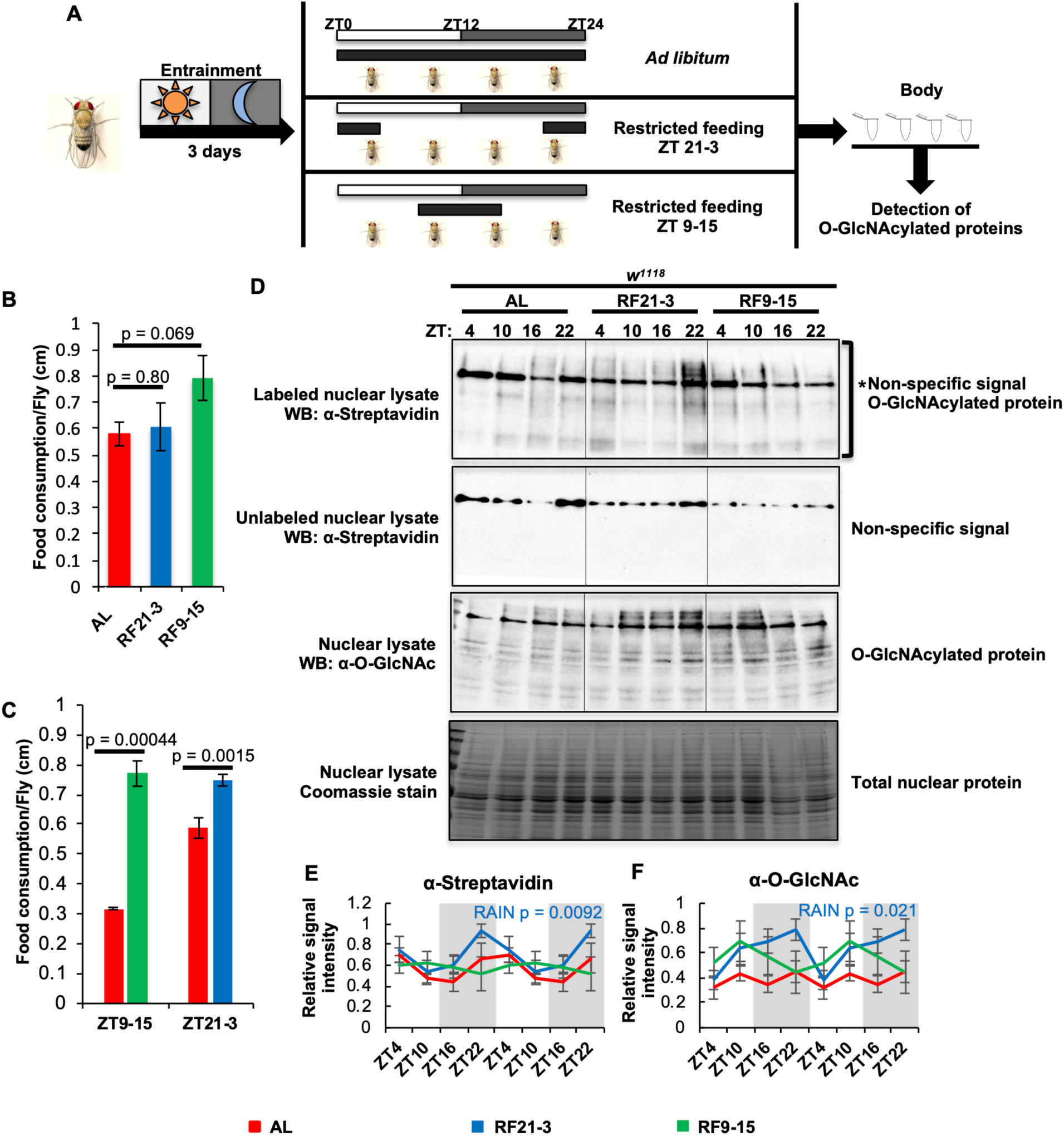
Time-restricted feeding (TRF) at natural feeding time strengthens the rhythm of protein O-GlcNAcylation. (A) Schematic showing the study design. Wild type (*w^1118^*) flies were first entrained in *ad libitum* (AL) conditions for 3 days in 12hr light:12hr dark (LD) cycles and then divided into 3 groups. The AL group continued to have food available all the time, the natural feeding (RF21-3) group was only fed between ZT21-3, and the mistimed feeding group (RF9-15) was fed during ZT9-15. The black bars indicate the time when food was available. Body tissues were collected after 6 days of feeding treatment and subjected to O-GlcNAcylation analysis. (B) Food consumption of AL, RF21-3 and RF9-15 groups over a 24-hr period measured using CAFE assay (n=4, in comparison to AL group, RF21-3 p=0.80 and RF9-15 p=0.069, two-tailed Student’s t-test). (C) CAFE assay to compare food consumption of TRF groups to that of AL group during the indicated time window (n=3, in comparison to AL group, RF21-3 p=0.0015 and RF9-15 p=0.00044, two-tailed Student’s t-test). (D) Western blots showing daily rhythms in O-GlcNAcylation in flies fed AL, between ZT21-3, or ZT9-15. O-GlcNAcylation was detected using two methods: (top 2 panels) chemoenzymatic labeling in combination with immunoblotting with *α*-streptavidin and (third panel) immunoblotting with *α*-O-GlcNAc. Unlabeled samples (second panel) were processed in parallel to labeled samples (top panel) to identify non-specific signal. Total nuclear proteins stained by Coomassie blue (bottom) were used for normalization. The asterisk (top panel) indicates non-specific signal. (E, F) Quantification of protein O-GlcNAcylation detected using enzymatic labeling (n=4; AL: p=0.045, RF21-3: p=0.0092, RF9-15: p=0.83, RAIN; AL vs RF21-3: mesor p=0.05, amplitude p=0.62, phase p=0.3, CircaCompare) and *α*-O-GlcNAc immunoblotting (n=4; AL: p=0.83, RF21-3: p=0.021, RF9-15: p=0.69; RAIN). Data are double plotted, normalized (peak=1), and presented as mean ± SEM. The biological replicates of AL group that are quantified in panel (E) are the same four replicates quantified in Fig. 1B. In panel (D), a different biological replicate of AL group, other than the one shown in Fig 1A, was ran together on the same gel with RF21-3 and RF9-15 groups.

The daily O-GlcNAcylation profile of nuclear proteins in WT body tissue was determined using the chemoenzymatic labeling method as described above. In the RF21-3 group, the oscillation of protein O-GlcNAcylation was strengthened, as indicated by comparing RAIN statistics of AL and RF21-3 groups (Fig. 3D top panel and E). The RF9-15 group, however, exhibited a severely dampened O-GlcNAcylation rhythm despite having access to food for the same duration of time and consuming roughly the same amount as the RF21-3 group (Fig. 3D top panel and E). These results derived from quantification of biotin-conjugated O-GlcNAcylated proteins detected using *α*-streptavidin were further confirmed by western blots using an *α*-O-GlcNAc antibody (Fig. 3D third panel from top and F). On one hand, results from TRF experiments support our hypothesis that feeding-fasting rhythm drives protein O-GlcNAcylation rhythm as consolidated feeding activity in the RF21-3 group led to more robust nuclear O-GlcNAcylation rhythm. On the other hand, since TRF at unnatural feeding time (RF 9-15) did not simply produce a phase shift of protein O-GlcNAcylation rhythm, we postulate that the circadian clock must play additional role(s) in regulating protein O-GlcNAcylation rhythm besides the regulation of feeding-fasting cycles.

### Circadian clock regulates the GFAT-catalyzed rate limiting step in HBP

Combing through CirGRDB (Li et al., 2018), a mammalian circadian transcriptomic database, we found that all the HBP enzymes have been identified as rhythmic genes in the mouse liver in at least one study, even though the phase of some transcripts may be variable between studies (Table S1) (Hughes et al., 2009; Hughes et al., 2012; Jouffe et al., 2013; Du et al., 2014; Janich et al., 2015). Moreover, previous proteomic or phosphoproteomic analyses indicated that some of these HBP enzymes exhibit rhythmic protein expression or phosphorylation status (Mauvoisin et al., 2014; Robles et al., 2017). We therefore reasoned that the circadian clock may affect one or more of the HBP enzymes to regulate protein O-GlcNAcylation rhythm. To identify key HBP effector(s) of the circadian clock, we performed targeted metabolomic analysis using hydrophilic interaction liquid chromatography coupled to mass spectrometry (HILIC-MS/MS) on body tissues of WT flies subjected to TRF between ZT21-3 and ZT9-15. We reasoned that measurements of HBP metabolites in combination with TRF treatments will allow us to evaluate potential time-of-day differences in HBP enzyme activities and identify key clock-controlled mechanism(s) that facilitate rhythmic protein O-GlcNAcylation when food is consumed at natural feeding time. These mechanisms are also expected to limit excessive O-GlcNAcylation when food intake occurs during non-optimal time during the day-night cycle. Standard curves of HBP metabolites were generated to obtain more reliable concentration measurements for each metabolite (Fig. S3A). We were able to detect 8 to 9 out of 13 metabolites in the HBP, but were not able to differentiate between GlcNAc-6-P and GlcNAc-1-P.

In both TRF groups, we observed robust cycling of all HBP metabolites (RAIN p<0.001), except for glucosamine and glucosamine-6-phosphate. The cycling HBP metabolites were found to peak after the respective feeding time in both TRF groups, suggesting that metabolite levels are significantly elevated in response to nutrient influx (Fig. 4A). We explored whether there was a significant difference in daily rhythmicity between HBP metabolites in the two TRF groups and found that fructose-6-phosphate is the most statistically different metabolite between the two groups (Dataset S3). In comparison to RF21-3 flies, the RF9-15 group showed a significant accumulation of fructose-6-phosphate right after the time of feeding, indicating that fructose-6-phosphate was not metabolized to glucosamine-6-phosphate by glutamine-fructose-6-phosphate amidotransferase (GFAT) (Fig. 2D and 4A). This would explain the lower UDP-GlcNAc peak level in the RF9-15 group (at ZT 24) as compared to that in the RF21-3 group (at ZT12) (Fig. 4A; t-test p=0.038). Our results therefore suggest that GFAT, the rate limiting enzyme in the HBP (Hart et al., 2011; Mishra et al., 2011; Yang and Qian, 2017; Hart, 2019), represents a key clock-controlled regulator that inhibits metabolic flow into the HBP during mistimed feeding.

**Figure 4.**
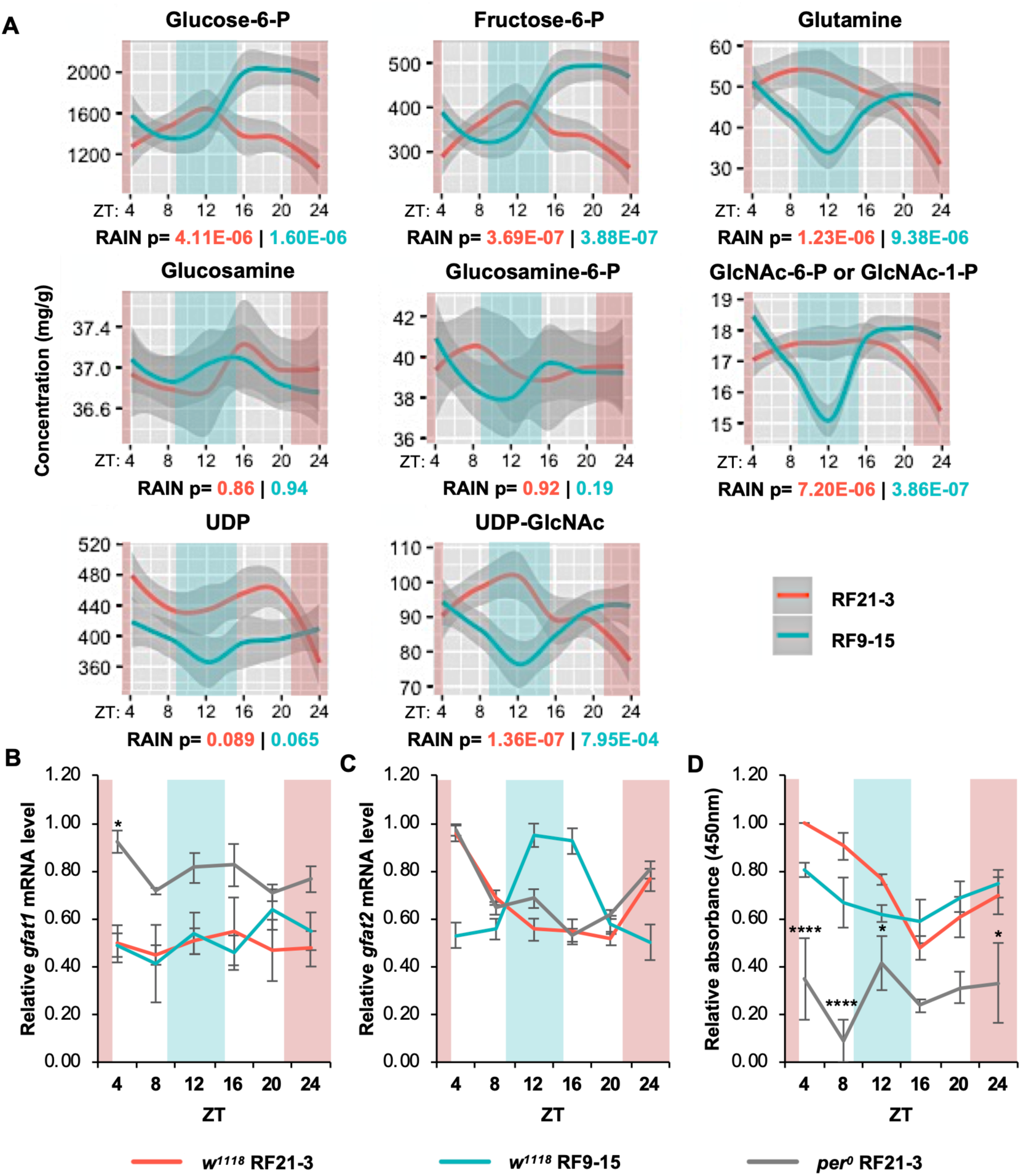
Glutamine-fructose-6-phosphate aminotransferase (GFAT) is a key enzyme that is regulated by circadian clock. (A) Line graphs showing daily rhythms of HBP metabolites detected by targeted metabolomics in WT (*w^1118^*) flies fed between ZT21-3 (red) or ZT9-15 (blue) (n=6; p values from RAIN analysis are presented under each graph). Gray-shaded regions represent 95% confidence interval. (B, C) Expression of *gfat1* and *gfat2* mRNA in three groups of flies: WT RF21-3, *per^0^* RF21-3, and WT RF9-15. (n=3; *gfat1* mRNA: WT RF21-3: p=0.73, WT RF9-15: p=0.59, *per^0^* RF21-3: p=0.76; *gfat2* mRNA: WT RF21-3: p=1.58E-05, WT RF9-15: p=0.00019, *per^0^* RF21-3: p=3.89E-06, RAIN); (*gfat2* mRNA: WT vs *per^0^* RF21-3: mesor p=0.23, amplitude p=0.53, phase p=0.66; WT RF21-3 vs RF9-15: mesor p=0.95, amplitude p=0.40, phase p=1.12E-12; CircaCompare). (D) Line graph representing daily GFAT enzymatic activity in WT (*w^1118^*) flies fed between ZT21-3 or ZT9-15 and *per^0^* flies fed between ZT21-3 (n=3; WT RF21-3: p=5.73E-07, WT RF9-15: p=0.041, *per^0^* RF21-3: p=0.92, RAIN; WT RF21-3 vs RF9-15: mesor p=0.14, amplitude p=0.01, phase p=0.027, CircaCompare). (B-D) Data are presented as mean ± SEM. Colored shading denotes the feeding periods for TRF treatment groups, RF21-3 (red) and RF9-15 (blue). The asterisks denote the comparison between WT RF21-3 and *per^0^* RF21-3 using two-way ANOVA with *post-hoc* Tukey’s HSD tests. *p<0.05, ****p<0.0001.

### Clock-controlled post-transcriptional mechanism imposes a stronger effect on GFAT enzymatic activity than feeding-dependent *gfat* mRNA expression

As the key molecular components of circadian clock are all transcription factors (Glossop et al., 1999; Hardin, 2011), we first investigated whether *gfat* is transcriptionally regulated by the circadian clock and/or feeding-fasting cycle. We assayed the mRNA levels of *gfat1* and *gfat2*, two functionally equivalent paralogues, in body tissues of WT RF21-3, WT RF9-15 and *per^0^* RF21-3 groups. *gfat1* mRNA did not exhibit daily rhythmicity in all three groups (Fig. 4B). On the other hand, we observed robust oscillation of *gfat2* mRNA levels with a peak right after the respective feeding window in all three groups (Fig. 4C). Interestingly, comparison of WT and *per^0^* RF21-3 flies revealed very similar daily *gfat2* mRNA profiles, suggesting that feeding time, rather than the molecular clock, is the key determinant in regulating rhythmic *gfat2* mRNA expression (Fig. 4C). Since the expression of *gfat2* is 5 to 10-fold higher than that of *gfat1* (Fig. S3B-D), we speculate that *gfat2* is the major paralogue that contributes to GFAT enzyme activity in fly bodies.

Since GFAT phosphorylation has been shown to oscillate in mouse liver over a 24-hour period (Robles et al., 2017), we hypothesized that the circadian clock may also modulate fly GFAT enzymatic activity by post-transcriptional regulation. To test this hypothesis, we assayed GFAT activity in the same three groups of flies described above over a 24-hr cycle. When WT flies were subjected to TRF at natural feeding time (RF21-3), we observed robust oscillation of GFAT enzymatic activity in whole body tissues that peaks immediately after the time of feeding (Fig. 4D). In comparison, GFAT activity in *per^0^* flies fed in the same time window showed significantly lower GFAT enzyme activity with no apparent daily oscillation despite similar patterns of *gfat2* mRNA expression (Fig. 4C). This suggests a vital role for the circadian clock in driving daily rhythms of GFAT activity through post-transcriptional regulation. Finally, we observed that the time of feeding can also modulate GFAT enzymatic activity rhythm; WT flies subjected to TRF at unnatural feeding time (RF9-15) exhibited dampened GFAT activity rhythm that has a phase difference of −3.48 as compared to that observed for the WT RF21-3 group (Fig. 4D). This could be attributed to the phase shift in *gfat2* mRNA rhythm that likely led to changes in daily expression of GFAT protein. Taken together, our results indicate that food-dependent *gfat2* mRNA induction and clock-controlled post-transcriptional mechanism collaborate to drive robust daily oscillation of GFAT activity.

### Rhythmic protein O-GlcNAcylation was not observed in *per^0^* clock mutant despite time-restricted feeding at natural feeding time

If GFAT activity rhythm is strongly regulated by clock-controlled post-transcriptional mechanism(s), it is conceivable that arrhythmic clock mutant flies would exhibit impaired O-GlcNAcylation even if fed at natural feeding time. We detected nuclear protein O-GlcNAcylation in body tissues of WT and *per^0^* flies under RF21-3 condition using chemoenzymatic labeling and western blotting by *α*-O-GlcNAc antibody. In comparison to WT, nuclear protein O-GlcNAcylation of *per^0^* flies did not exhibit clear daily rhythmicity even though they were fed during natural feeding time (Fig. 5A-C). Since results from CAFE assay confirmed that WT and *per^0^* flies consumed similar amount of food under TRF treatment (Fig. 5D, t-test p=0.35), we ruled out the possibility that the altered O-GlcNAcylation rhythm in *per^0^* flies was due to different amount of food consumption. Our results therefore support that clock-controlled post-transcriptional mechanism represents the primary buffering mechanism to inhibit excessive O-GlcNAcylation when animals feed at unnatural time.

**Figure 5.**
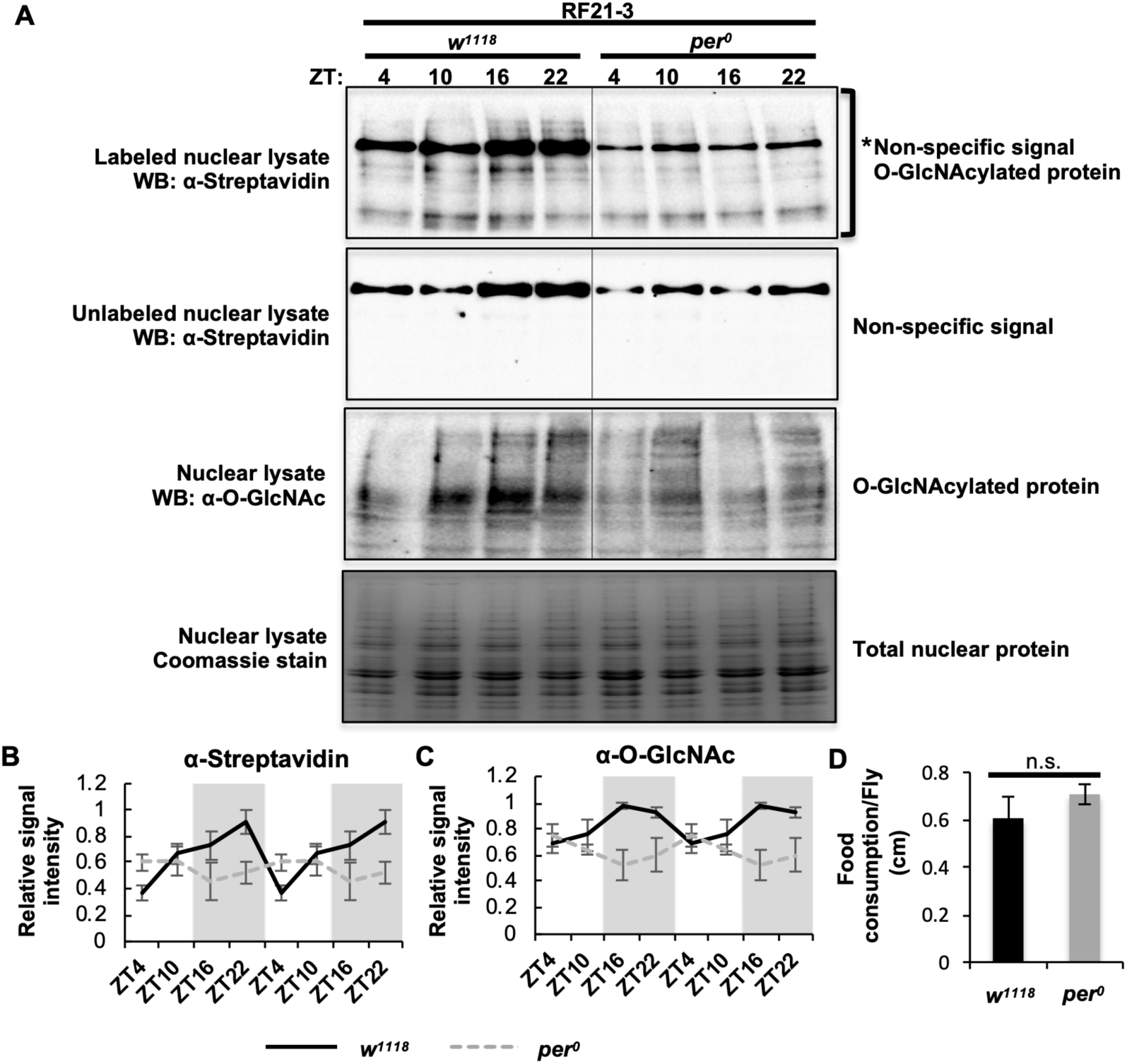
The circadian clock regulates daily rhythms in protein O-GlcNAcylation through multiple mechanisms. (A) Western blots showing daily rhythms in nuclear protein O- GlcNAcylation in WT (*w^1118^*) and *per^0^* flies fed between ZT21-3 (natural feeding time). Protein O-GlcNAcylation was detected using two methods: (top 2 panels) chemoenzymatic labeling in combination with immunoblotting with *α*-streptavidin and (third panel) immunoblotting with *α*-O-GlcNAc. Unlabeled samples (second panel) were processed in parallel to labeled samples (top panel) to identify non-specific signal. Total nuclear proteins stained by Coomassie blue (bottom) were used for normalization. The asterisk (top panel) denotes non-specific signal. (B, C) Quantification of protein O-GlcNAcylation detected using enzymatic labeling (n=4; WT: p=0.0096, *per^0^*: p=0.44, RAIN; WT vs *per^0^*: p=0.011, DODR), and using *α*-O-GlcNAc (n=4; WT: p=0.015, *per^0^*: p=0.28, RAIN; WT vs *per^0^*: p=0.017, DODR). Data are double plotted. Gray shading indicates the dark period. (D) Food consumption of WT and *per^0^* flies over 24-hr period as measured by CAFE assay (n=4; p=0.35; two-tailed Student’s t-test). Data are presented as mean ± SEM.

### Mathematical modeling identifies timing of food intake and presence of an intact molecular clock as key factors that produce daily rhythms of protein O-GlcNAcylation

To assess the extent to which feeding time and clock-controlled mechanisms contribute to protein O-GlcNAcylation rhythm, we formulated a mathematical model by simulating the results of WT flies subjected to TRF at natural feeding time (ZT21-3) (See Table 1 for equations and variables). In WT flies, the circadian clock drives feeding-fasting cycles to provide timely nutrient influx, which transiently induces the expression of *gfat2* and promotes GFAT enzymatic activity (Fig. 6A). Additionally, there is(are) clock-controlled mechanism(s) at the post-transcriptional level to stimulate GFAT enzyme activity after food intake. GFAT, the rate limiting enzyme of HBP, increases the HBP metabolic flow to produce the end product, UDP-GlcNAc. As the donor substrate of protein O-GlcNAcylation, UDP-GlcNAc feeds into the activity of O-GlcNAc transferase (OGT) and increases O-GlcNAcylation to peak level. Subsequently, O-GlcNAcase (OGA) removes the O-GlcNAc groups on proteins to reset the O-GlcNAcylation status every day (Fig. 6A).

**Figure 6.**
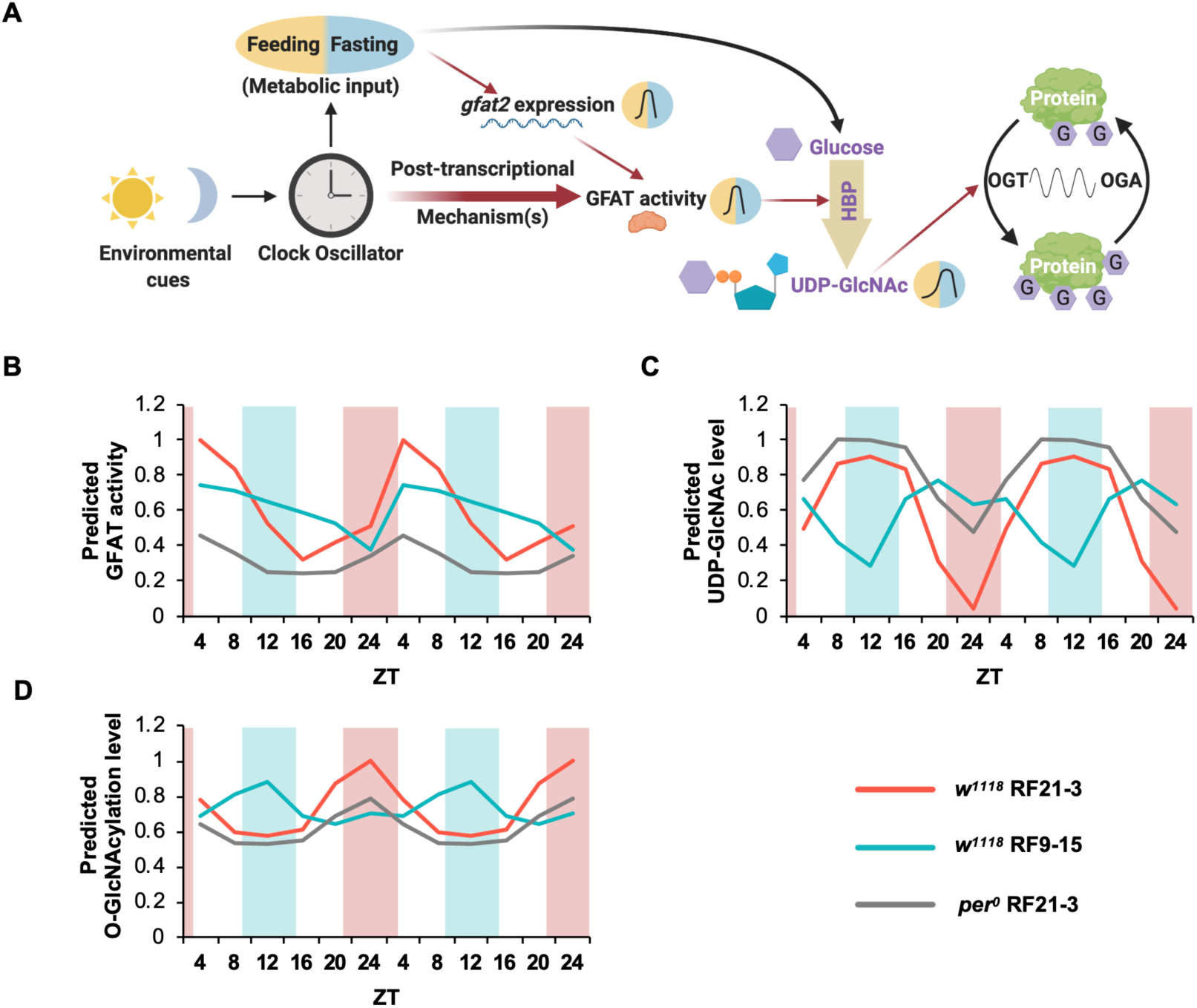
A mathematical model describes and predicts daily protein O-GlcNAcylation rhythm based on timing of food intake and molecular clock function. (A) Schematic model showing the regulation of protein O-GlcNAcylation by the circadian clock through feeding activity and post-transcriptional mechanism(s). Yellow background indicates feeding period while the blue background indicates fasting period. Red arrows denote activation, and the thicker red arrows denote a stronger effect. (B-D) Line graphs showing daily rhythms of (B) GFAT activity, (C) UDP-GlcNAc, and (D) protein O-GlcNAcylation levels as predicted using the mathematical model (Table 1). Time of feeding for WT RF21-3, WT RF9-15 and *per^0^* RF21-3 fly groups were used as input for the model and the circadian clock is assumed to be functional in WT but not in *per^0^* flies. Colored shading denotes the feeding periods for TRF treatment groups, RF21-3 (red) and RF9-15 (blue).

**Table 1.** Mathematical model describing key parameters that regulate daily rhythms of protein O-GlcNAcylation

| Variables | Description |
| --- | --- |
| $F_t$ | Daily feeding pattern of flies |
| $G2_t$ | Daily *gfat2* mRNA level |
| $C_t$ | Clock-controlled post-transcriptional mechanism(s) |
| $a$ | Parameter to indicate whether flies have an intact circadian clock. WT flies ($w^{1118}$): $a=1$; clock mutant flies ($per^0$): $a=0$ |
| $G_t$ | Daily GFAT activity level |
| $U1_t$ | Level of UDP-GlcNAc calculated using GFAT activity |
| $U2_t$ | Level of UDP-GlcNAc calculated using feeding activity |
| $U_t$ | Overall daily UDP-GlcNAc levels |
| $O_t$ | Daily O-GlcNAcylation levels |

To validate this model, we input the appropriate parameters for the WT RF9-15 group and predicted daily rhythms in GFAT activity, UDP-GlcNAc, and protein O-GlcNAcylation levels (Fig. 6B-D, S4B and S5A). In agreement with our experimental results, the model predicted similar phase but dampened GFAT activity rhythm for the WT RF9-15 group as compared to the WT RF21-3 group (Fig. 6B and S5A). In addition, the model predicted lower amplitude of UDP-GlcNAc rhythm that peaks at the opposite phase, consistent with the fact that feeding significantly impacts metabolite levels (Fig. 4A, 6C and S5A), and dampened O-GlcNAcylation rhythm with a slight peak around ZT12 (Fig. 3E-F, 6D and S5A).

We also validated our model by comparing the predictions to the experimental data obtained for *per^0^* RF21-3 flies (Fig. S4C and S5B). Without a functional molecular clock, our model predicted that the daily rhythms of GFAT activity, UDP-GlcNAc, and protein O-GlcNAcylation levels would all be dampened, thus recapitulating our experimental data (Fig. 5A-C, 6B-D and S5B). In summary, although there are minor differences between the predicted rhythms from the model and our experimental observations, our mathematical model successfully predicted the phase and amplitude of daily GFAT activity rhythms and rhythms of UDP-GlcNAc and protein O-GlcNAcylation levels in WT RF9-15 and *per^0^* RF21-3 groups. This suggests that our model included the major parameters that regulate daily protein O-GlcNAcylation rhythm.

## DISCUSSION

In this study, we sought to investigate the mechanisms by which nutrient influx through clock-controlled feeding-fasting cycles regulate time-of-day-specific protein O-GlcNAcylation and cellular protein functions. Despite the important roles of O-GlcNAcylation in maintaining cellular homeostasis, most studies on protein O-GlcNAcylation were either conducted in tissue culture system (Cheung and Hart, 2008; Taylor et al., 2008; Taylor et al., 2009) or in metabolic disease models (Yang et al., 2008; Bond and Hanover, 2013; Perez-Cervera et al., 2013; Ruan et al., 2014; Ruan et al., 2017; Hart, 2019; Nie and Yi, 2019), and failed to take into account the intrinsic importance of O-GlcNAc cycling in the circadian timeframe. Here, we report daily rhythms of protein O-GlcNAcylation under normal physiological conditions by sampling WT fly tissues over a 24-hr period. Under *ad libitum* conditions, we established strong correlations between daily rhythms in food intake, UDP-GlcNAc, and cellular protein O-GlcNAcylation level. Despite the fact that O-GlcNAcylation is known to be nutrient sensitive, this is the first study to establish a clear relationship between feeding-fasting cycle and O-GlcNAcylation rhythm in whole animals. Furthermore, by subjecting flies to TRF, we uncovered a clock-controlled buffering mechanism that inhibits excessive O-GlcNAcylation when animals feed at non-optimal times in the day-night cycle. Specifically, we showed that the activity of GFAT, the rate limiting enzyme of HBP, mediates this buffering mechanism. Our results indicate that although both feeding-induced *gfat2* mRNA expression and clock-controlled post-transcriptional regulation of GFAT activity contribute to the regulation of GFAT activity, the latter appears to have greater impact on the daily oscillation of GFAT activity, which contributes to rhythms in UDP-GlcNAc production and protein O-GlcNAcylation level.

To describe key factors that regulate daily O-GlcNAcylation rhythm and identify potential parameters we have not yet considered, we generated a mathematical model that parameterized timing of food intake and presence of an intact molecular clock to predict protein O-GlcNAcylation rhythms (Table 1). Our model correctly predicted the dampened daily O-GlcNAcylation rhythms in WT RF9-15 and *per^0^* RF21-3 treatment groups (Fig. 6D), suggesting that the parameters in our model include major factors that impact O-GlcNAcylation rhythms. Nevertheless, the O-GlcNAcylation rhythm predicted in WT RF9-15 and *per^0^* RF21-3 groups still showed weak rhythmicity, which is somewhat different from our experimental results (Fig. 3D-F and Fig. 5A-C). There are two possible reasons for the minor discrepancy: (1) The variability in experimental results makes it difficult to conclude whether the O-GlcNAcylation is still weakly rhythmic or totally arrhythmic in these flies (compare Fig. 3E-F); (2) Our model still lacks some parameters that modulate protein O-GlcNAcylation rhythms. For example, the activity of OGT has been reported to be regulated by GSK3β (Kaasik et al., 2013), insulin receptor (Whelan et al., 2008), AMPK (Bullen et al, 2014; Xu et al., 2014) and calcium/calmodulin-dependent protein kinase II (CaMKII) (Ruan et al., 2017). All of these kinases are sensitive to cellular metabolic status and are potentially regulated by either circadian clock or food intake. Additionally, both OGT and OGA are O-GlcNAcylated (Tai et al., 2004; Khidekel et al., 2007; Seo et al., 2016), suggesting that daily O-GlcNAcylation rhythm might modulate the temporal activity of its own writer and eraser. Future investigations on circadian OGT and OGA activity might shed light on the regulation of O-GlcNAcylation rhythm and further improve our mathematical model.

Future study to investigate how GFAT activity is regulated by clock-controlled post-transcriptional mechanism(s) is also warranted. Our results showed that GFAT activity is regulated by both feeding-induced gene expression and clock-controlled post-transcriptional mechanism(s). GFAT activity has been shown to be regulated by PKA- and AMPK-directed phosphorylation *in vitro* (Zhou et al., 1998; Chang et al., 2000; Graack et al., 2001; Hu et al., 2004; Eguchi et al., 2009). Interestingly, in mouse hepatic tissue, one of the previously identified PKA-directed phosphorylation sites on GFAT1, which activates its activity, was found to exhibit daily oscillation that peaks slightly after feeding phase (Zhou et al., 1998; Robles et al., 2017). This evidence supports our finding that GFAT activity peaks after natural feeding time in WT flies. We expect that future investigation on GFAT phosphorylation or other post-transcriptional mechanism will inform on the circadian regulation of GFAT activity.

Despite our findings that nuclear proteins extracted from fly body tissues exhibit pronounced daily rhythms in O-GlcNAcylation, the identity of cellular proteins that are regulated in this manner will need to be uncovered in circadian O-GlcNAc proteomic analysis. Nevertheless, given the diverse cellular proteins that are known to be regulated by O-GlcNAcylation, a significant portion of the proteome may be regulated in this manner. As explained earlier, this study focused primarily on O-GlcNAcylation analysis of total nuclear proteins. One of the major functions of nuclear proteins is transcription. Indeed, O-GlcNAcylation is known to modulate almost every step of transcription; the activities of transcriptional regulators including RNA polymerase II, histones, TATA-binding protein, and ten eleven translocation (TET) enzymes, are all heavily regulated by O-GlcNAcylation (Kelly et al., 1993; Fujiki et al., 2011; Ranuncolo et al., 2012; Lewis et al., 2016; Bauer et al., 2015; Wang et al., 2015; Hrit et al., 2018; Hardivillé et al., 2020). Moreover, O-GlcNAcylation is also known to be enriched in nuclear pore complexes, although the specific function of O-GlcNAcylation in nucleocytoplasmic trafficking is still unclear (Ruba and Yang, 2016). O-GlcNAcylation is also essential for safeguarding the functions of multiple biological processes in the cytoplasm. Translational factors, ribosomal proteins, proteins in mitochondrial respiration chain, cytoskeletal proteins are all O-GlcNAcylated (Holt et al., 1987; Datta et al., 1989; Slawson et al., 2005; Hu et al., 2009; Zeidan et al., 2010; Hart, 2019). The prevalence of cellular protein O-GlcNAcylation suggests that it could represent a key mechanism by which organisms leveraged to reinforce the robustness of circadian physiology. We expect that future examination on cytoplasmic protein O-GlcNAcylation rhythm as well as circadian O-GlcNAc proteomics to identify substrate proteins would certainly reveal the physiological processes that are regulated in this manner.

Although most of this study focused on O-GlcNAcylation of proteins extracted from fly bodies as body tissues are known to be more metabolically sensitive in flies, it is interesting to note that a small portion of metabolites also oscillate in fly head tissues in our GC-MS data. Given that O-GlcNAcylation of PER and CLOCK was shown to exhibit daily rhythms in fly heads (Kaasik et al., 2013; Li et al., 2019), we expect that a significant number of fly brain proteins may also be sensitive to nutrient signals and could be regulated in a phase-specific manner by daily O-GlcNAcylation rhythm.

Since O-GlcNAcylation has been described in a wide range of species including *Caenorhabditis elegans*, fruit flies, mice, and humans (Olszewski et al., 2010; Levine and Walker, 2016), we expect that O-GlcNAcylation could represent a conserved molecular mechanism that integrates circadian and metabolic signals in animals. To investigate whether UDP-GlcNAc level also exhibits daily rhythmicity in the mouse model and could potentially drive rhythms in protein O-GlcNAcylation, we searched published circadian metabolomics datasets. We found that most of the studies were not able to detect UDP-GlcNAc, perhaps due to the use of different platforms and metabolomics methods (Hatori et al., 2012; Eckel-Mahan et al., 2012; Eckel-Mahan et al., 2013; Chaix et al., 2014; Masri et al., 2014; Abbondante et al., 2016; Masri et al., 2016; Krishnaiah et al., 2017). Among the two studies that detected UDP-GlcNAc, one showed that TRF of clock mutant mice on high fat diet (HFD) was sufficient to drive daily oscillation of UDP-GlcNAc in liver (Chaix et al., 2018), while the other study failed to detect rhythmic UDP-GlcNAc level in brown adipose tissues under *ad libitum* condition (Dyar et al., 2018). Based on these limited results, it is clear that at least some mammalian tissues would display daily rhythms in UDP-GlcNAc. Whether these rhythms in fact translate to rhythms in protein O-GlcNAcylation, as we observed in fly tissues, will need to be confirmed in future experiments.

In summary, we showed that HBP integrates circadian and metabolic signals to drive daily rhythms in protein O-GlcNAcylation. Since we and others have previously reported that global manipulation of OGT and OGA activity and disruption of clock protein O-GlcNAcylation can alter circadian rhythms in whole animals (Kim et al., 2012; Kaasik et al., 2013; Li et al., 2013; Li et al., 2019), it is conceivable that daily O-GlcNAc cycling represents a key post-translational mechanism that regulates circadian physiology. Future investigations that examine functional consequences of protein O-GlcNAcylation and the interplay between O-GlcNAcylation and other PTMs will reveal how daily rhythms in O-GlcNAcylation manifest into rhythms in circadian physiology. Our results will facilitate future efforts to integrate key metabolic signaling pathways to obtain a systems-level understanding of metabolic regulation of circadian physiology. Finally, this study could shed light on the mechanisms by which lifestyle and eating habits common in modern society, especially night-shifted eating, contribute to the current epidemic of metabolic disorders and obesity as the misalignment of human lifestyles and natural day-night cycles continues to increase.

## MATERIALS AND METHODS

### Animals

*w^1118^*; UAS-FLAG-*ogt* flies were described in Li et al. (2019).

### Manipulation of metabolic input with time-restricted feeding

0-4*^-^*day old flies were entrained in 12 hr light: 12 hr dark (LD) cycles at 25°C for three days and split into three groups: *ad libitum* (AL), RF21-3 and RF9-15 groups. RF21-3 and RF9-15 groups were fed at ZT21-3 and ZT9-15 respectively for six days. During the fasting period, flies were kept on 2% agar as water source. Flies were collected for O-GlcNAcylation analysis at the conclusion of the 6-day feeding treatments.

### CApillary FEeder (CAFE) assay to measure daily rhythms of fly feeding

Daily feeding rhythms of flies fed *ad libitum* (AL) were measured by CAFE assay (Ja et al., 2007; Xu et al., 2008) after three days of entrainment in LD cycle. Food was provided to flies (groups of ten flies per biological replicate) as 5% sucrose solution in calibrated pipettes during a 24-hour training period. Pipettes were replaced after the training period, and food consumption was measured at ZT 4, 10, 16, 22 for 2-hour periods for two consecutive days. The evaporation rate was measured using pipettes without flies feeding on them. CAFE assay was also performed with TRF treatments, RF21-3 and RF 9-15. Just as for AL feeding, flies were fed with 5% sucrose solution in calibrated pipettes during the feeding period (ZT21-3 or ZT9-15) and changed into pipettes filled with water during the fasting period. One-day food consumption was measured at the conclusion of their feeding period.

### Chemoenzymatic labeling of O-GlcNAcylated proteins

Nuclear proteins from fly bodies were extracted as previously described (Kwok et al., 2015; Li et al., 2019) with modifications. Fly bodies were grinded in liquid nitrogen using mortar and pestles and resuspended in lysis buffer (20mM HEPES pH7.5, 1mM DTT, Roche EDTA-free protease inhibitor cocktail). After incubating for 15 mins, a glass dounce homogenizer was used to disrupt the cell membrane. Samples were passed through cell strainers by centrifugation at 300 g for 1 min at 4°C. Supernatant was collected and nuclei were pelleted by centrifuging at 6700 rpm for 15 min at 4°C. The supernatant was collected as cytoplasmic fraction. The crude nuclear pellet was rinsed in wash solution (20mM HEPES pH7.5, 10% glycerol, 150mM NaCl, 0.1% TritonX-100, 1mM DTT, 1mM MgCl_2_, 0.5mM EDTA, 10mM NaF, Roche protease inhibitor cocktail) twice, resuspended in nuclear extraction buffer (20mM HEPES pH7.5, 10% glycerol, 350mM NaCl, 0.1% TritonX-100, 1mM DTT, 1mM MgCl_2_, 0.5mM EDTA, 10mM NaF, Roche protease inhibitor cocktail) and incubated for 30 min at 4°C on a rotator. After spinning at 13000 rpm for 15min at 4°C, the supernatant was collected as nuclear fraction. 50 *μ*g nuclear protein was precipitated using chloroform-methanol precipitation and dissolved in 40 *μ*l 1% SDS with 20mM HEPES pH7.9 by heating at 95°C for 5-10 min. Samples were azide-labeled using Click-iT O-GlcNAc Enzymatic Labeling System and reacted with biotin alkyne using Click-IT Biotin Protein Analysis Detection Kit (Thermo Fisher Scientific, Waltham, MA) following manufacturer’s protocol. Unlabeled samples were processed in parallel for background deduction; the azide-labeling enzyme, GalT1 (Y289L), was excluded in these reactions.

### Western blotting of protein extracts and protein quantification

Western blot was performed after chemoenzymatic labeling to detect O-GlcNAcylated proteins using α-streptavidin-HRP (Cell Signaling Technologies, Danvers, MA) (1:3000). O-GlcNAcylated protein was normalized to total proteins stained with Coomassie blue (Bio-Rad, Hercules, CA) and the normalized signal intensity was presented as a proportion of the peak O-GlcNAcylation intensity (Peak=1). To confirm the results based on chemoenzymatic labeling, we also used α-O-GlcNAc (Abcam, Cambridge, MA) (1:1000) to detect O-GlcNAcylation in nuclear lysates. The O-GlcNAc antibody was validated using flies overexpressing OGT (Li et al., 2019). Our results indicated that there are higher levels of protein O-GlcNAcylation in the OGT overexpressor flies especially for proteins ranging from 37kD to 250kD (Fig. S6A). For this reason, only proteins within this size range were included in our quantification. Coomassie staining was performed to indicate equal amount of proteins in the labeling reactions and for normalization. The quality of nuclear and cytoplasmic extraction was validated by western blotting by α-Histone 3 (Sigma, St. Louis, MO) (1:2000) (Fig. S6B). Secondary antibodies used for western blotting are as follows: α-mouse-HRP (1:1000) (GE Healthcare, Piscataway, NJ) for α-O-GlcNAc primary antibody, α-rabbit-HRP (1:2000) (Sigma, St. Louis, MO) for α-H3 primary antibody.

### Sample preparation for metabolomics analysis

10 mg of tissue per sample were grinded using metal beads and extracted twice in 1 ml of methanol, chloroform and water (5:2:2) mixture for 20 min at 4 °C under constant agitation. After centrifugation at 14,000 g for 3 min, supernatants were pooled, evaporated to dryness under vacuum in a speedvac concentrator (Savant, ThermoFisher Scientific), and stored at −80 °C until GC-TOF MS or HILIC-TripleTOF MS/MS analysis.

### GC-TOF MS analysis

Samples were analyzed using a GC-TOF MS approach as previously described (Fiehn, 2016). Samples were randomized across the design using the MiniX database (Fiehn et al., 2005). A Gerstel MPS2 automatic liner exchange system (ALEX) was used to eliminate cross-contamination from sample matrix occurring between sample runs. 0.5 μl of sample was injected at 50°C (ramped to 250°C) in splitless mode with a 25 sec splitless time. An Agilent 6890 gas chromatograph (Agilent, Santa Clara, CA) was used with a 30 m long, 0.25 mm i.d. Rtx5Sil-MS column with 0.25 µm 5% diphenyl film; an additional 10 m integrated guard column was used (Restek, State College, PA). Chromatography was performed at a constant flow of 1 ml/min, ramping the oven temperature from 50°C for to 330°C over 22 mins. Mass spectrometry used a Leco Pegasus IV time of flight mass (TOF) spectrometer with 280°C transfer line temperature, electron ionization at −70 V and an ion source temperature of 250°C. Mass spectra were acquired from m/z 85–500 at 17 spectra/sec and 1750 V detector voltage.

Result files were exported to the servers and further processed by the metabolomics BinBase database (Kind et al., 2009). All database entries in BinBase were matched against the Fiehn mass spectral library of 1,200 authentic metabolite spectra using retention index and mass spectrum information or the NIST20 licensed commercial library (https://www.nist.gov/programs-projects/nist20-updates-nist-tandem-and-electron-ionization-spectral-libraries). Identified metabolites were reported if present in at least 50% of the samples per study design group (as defined in the MiniX database); output results were exported to the BinBase database and filtered by multiple parameters to exclude noisy or inconsistent peaks. Quantification was reported as peak height using the unique ion as default. Missing values were replaced using the raw data netCDF files from the quantification ion traces at the target retention times, subtracting local background noise. Sample-wise metabolite intensities were normalized by the total signal for all annotated analytes. Daily quality controls and standard plasma obtained from National Institute of Standards and Technology (NIST) were used to monitor instrument performance during the data acquisition. Metabolite data were normalized to the sum of intensity peaks of NIST plasma metabolites to avoid the effect of machine drift among each sample. Four samples were determined as outliers by comparing the total peak intensity of individual replicates at each time point. Replicates that exhibited 10-fold differences were removed from the dataset (Fig. S1B-C, Dataset S1). Data were then pareto scaled and heat maps were generated using Metaboanalyst (Xia et al., 2009; Chong et al., 2018). The metabolomics dataset has been deposited in the open metabolomics database, Metabolomics Workbench (https://www.metabolomicsworkbench.org/), under accession no. [ST001110]. Unknown metabolites can be visualized and investigated using the BinBase identifier (Dataset S1, column D) in https://binvestigate.fiehnlab.ucdavis.edu/#/.

### HILIC-TripleTOF MS/MS analysis

5 μl of resuspended sample was injected onto a Waters Acquity UPLC BEH Amide column (1.7 μm, 150 mm × 2.1 mm) with a Waters Acquity UPLC BEH Amide VanGuard pre-column (1.7 μm, 5 mm × 2.1 mm). Columns are maintained at 40°C. The mobile phases were prepared with two solutions: (A) 10 mM ammonium formate and 0.125% formic acid (Sigma, St. Louis, MO) in ultrapure water and (B) 10 mM ammonium formate and 0.125% formic acid (Sigma, St. Louis, MO) in 95:5 v/v acetonitrile : ultrapure water. Samples were eluted at 0.4 mL/min with the following gradient: 100% (B) at 0-2 min, 70% (B) at 2-7 min, 40% (B) at 7.7-9 min, 30% (B) at 9.5-10.25 min, 100% (B) at 10.25-12.75 min, 100% (B) until 16.75 min. Mass spectra were collected using an AB Sciex TripleTOF 6600 (SCIEX, Framingham, MA) in ESI (-) mode. Electron ionization voltage was −4.5 kV and spectra were acquired from m/z 60–1200 at 2 spectra/sec. Internal standard mix were injected along with each sample to monitor instrument performance during the data acquisition. A standard HBP metabolite mix, with the amount of each metabolites ranging from 6.25 ng to 50 ng, was used to generate the standard curve.

Mass spectra were deconvoluted, aligned and identified using MS-DIAL (Tsugawa et al., 2015). The metabolite data were normalized to the sum peak intensity of internal standard mix. Standard curves of the HBP metabolites were generated by plotting the normalized peak intensity against the amount of compound injected on the column. The amount of each HBP metabolites in fly samples was then calculated and converted to concentration (mg/g tissue) based on the injection volume and the input amount of fly tissues. The targeted metabolomics dataset will be deposited to Metabolomics Workbench and the accession number will be provided prior to publication.

### GFAT enzymatic activity assay

GFAT enzymatic activity was measured as described in Srinivasan et al. (2007) with the following modifications. Fly bodies were grinded in extraction buffer (60mM KH_2_PO_4_, pH7.8, 50mM KCl, 1mM EDTA, 1mM DTT) and sonicated to disrupt the cell membrane. Cell lysate was collected after centrifugation at 13000 rpm for 15 mins at 4°C. GFAT reactions were performed with 500 μg protein dissolved in reaction buffer (60mM KH_2_PO_4_, pH7.8, 50mM KCl, 1mM EDTA, 1mM DTT, 15mM fructose-6-P, 15mM L-glutamine). Two reactions were set up for each sample: positive reaction and negative reaction. In positive reaction, the endogenous GFAT catalyzed the reaction between fructose-6-P and glutamine to produce glutamate, which was the readout of GFAT activity, and reactions were incubated at 25°C for 90 mins and terminated by heating up at 95°C for 2 mins. In negative reaction, all the proteins in reaction, including GFAT, were denatured right away by heating up at 95°C for 2 mins and therefore, the original level of glutamate in fly lysate is measured for background deduction. After centrifugation at 13000 rpm for 15 mins, supernatant was collected and filtered using 10kD MWCO spin filters (Millipore Sigma, Burlington, MA) to deproteinate. The byproduct from GFAT catalyzed reaction, glutamate, was quantified using glutamate assay kit (Millipore Sigma) following the manufacturer’s protocol. 18μl of samples was used in each glutamate assay reaction. Absorbance was measured at 450nm using a TriStar LD 941 microplate reader (Berthold Technologies, Oak Ridge, TN). To determine the activity of GFAT, the absorbance of negative reactions was subtracted from that of positive reactions.

### Analysis of gene expression

Total RNA of fly body tissues was extracted using TRI reagent (Sigma, St. Louis, MO). Complementary DNA synthesis and real-time quantitative PCR were performed as previously described (Kwok et al., 2015; Li et al., 2019). Expression of target genes was normalized to non-cycling *cbp20* (Table S2).

### Statistical analysis

RAIN (Rhythmic Analysis Incorporating Nonparametric methods) (Thaben and Westermark, 2014) was used to determine rhythmicity, amplitude, phase and period length of metabolites, protein O-GlcNAcylation, steady state mRNA levels, and GFAT activity. Differences in daily rhythmicity were assessed using DODR (Detection of Differential Rhythmicity) (Thaben and Westermark, 2016) and differences in parameters (mesor, amplitude and phase) between two rhythms were analyzed using CircaCompare (https://rwparsons.shinyapps.io/circacompare/) (Parsons et al., 2020). RAIN and DODR were performed in R. Significance in differences between treatments were determined using GraphPad Prism 8.0 (GraphPad Software, La Jolla California USA). The difference of *gfat* mRNA or GFAT activity levels were assessed using two-way ANOVA with *post-hoc* Tukey’s HSD tests, while pair-wise comparisons between food consumption were analyzed using two-tailed Student’s t-test with equal variance. The coefficients for cross correlation between rhythms were calculated in R using “astsa” package (Shumway and Stoffer, 2017).

### Mathematical modeling

We generated a mathematical model to describe and predict O-GlcNAcylation rhythms using timing of fly feeding activity and the presence of molecular clock function. The model is designed based on Fig. 6A. We simulated the data of WT fly feeding activity at natural time (ZT21-3) using “astsa” package in R (Shumway and Stoffer, 2017), and calculated the rhythm of protein O-GlcNAcylation using the equations listed in Table 1. Description of variables are also included in Table 1. The input data for the model was the feeding activity of flies, which was first decomposed using “astsa” package to extract the rhythmic pattern (Fig. S4, Shumway and Stoffer, 2017). The decomposed feeding data was used to calculate the levels of *gfat2* mRNA. GFAT activity was determined based on both *gfat2* mRNA and clock-controlled post-transcriptional mechanism(s). The clock-controlled post-transcriptional mechanism(s) was(were) simulated to peak at the same time of maximum GFAT activity in RF21-3 flies. Using the daily GFAT activity and feeding activity, the UDP-GlcNAc rhythm was calculated. Finally, O-GlcNAcylation rhythm was determined based on UDP-GlcNAc levels. Data was presented as a proportion of the highest level of GFAT activity, UDP-GlcNAc, or O-GlcNAcylation (Highest value=1).

## ACKNOWLEDGMENTS

We would like to thank the West Coast Metabolomics Center at UC Davis for their technical support. This work is supported by National Institutes of Health grants R01 GM102225 and R01 DK124068 to JCC, and U24 DK097154 to the West Coast Metabolomics Center. XL is supported by the China Scholarship Council Fellowship, the UC Davis Jastro Graduate Research Fellowship, Marv Kinsey Scholarship, and Sean & Anne Duffey and Hugh & Geraldine Dingle Research Fellowship.

## AUTHOR CONTRIBUTIONS

XL, JCC, OF designed research; XL, IB, AC, TP, CAT, JJ performed research; XL, JCC, IB, OF, AC, YHL, JJ contributed to critical interpretation of the data. XL, JCC wrote the paper.

## RESOURCE AVAILABILITY

### Lead Contact

Further information and requests for resources and reagents should be directed to and will be fulfilled by the lead contact Joanna C. Chiu.

### Materials Availability

This study did not generate new unique reagents.

### Data and Code Availability

The untargeted metabolomics dataset has been deposited in the open metabolomics database, Metabolomics Workbench (https://www.metabolomicsworkbench.org/), under accession no. [ST001110]. The targeted metabolomics on HBP will be deposited to Metabolomics Workbench and the accession number will be provided prior to publication.

## SUPPLEMENTAL INFORMATION

**Figure S1.**
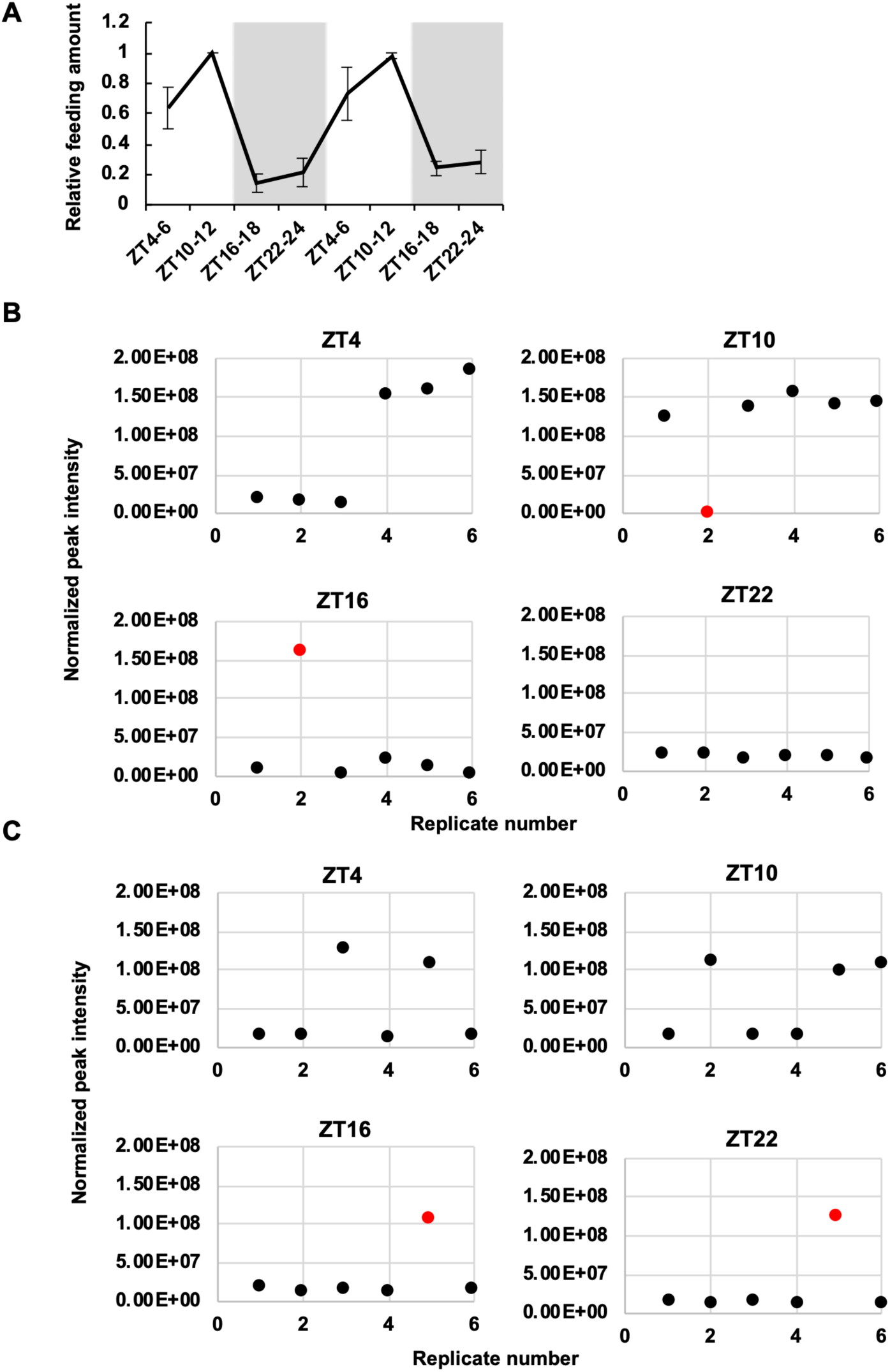
Analysis of fly feeding rhythms and circadian metabolomics. (A) Line graph showing the feeding-fasting cycles of WT (*w^1118^*) female flies over 2 day-night cycles (n=3, 10 flies per biological replicate; p=6.96E-07, RAIN). Data were normalized (peak=1) and presented as mean ± SEM. (B-C) Determination of outlier replicates in GC-MS metabolomics to be excluded from analysis. The quantity of the total metabolite peak intensity in each sample was graphed to identify outlier replicates in (B) body and (C) head samples. The red dots indicate outlier replicates. Related to Figure 2.

**Figure S2.**
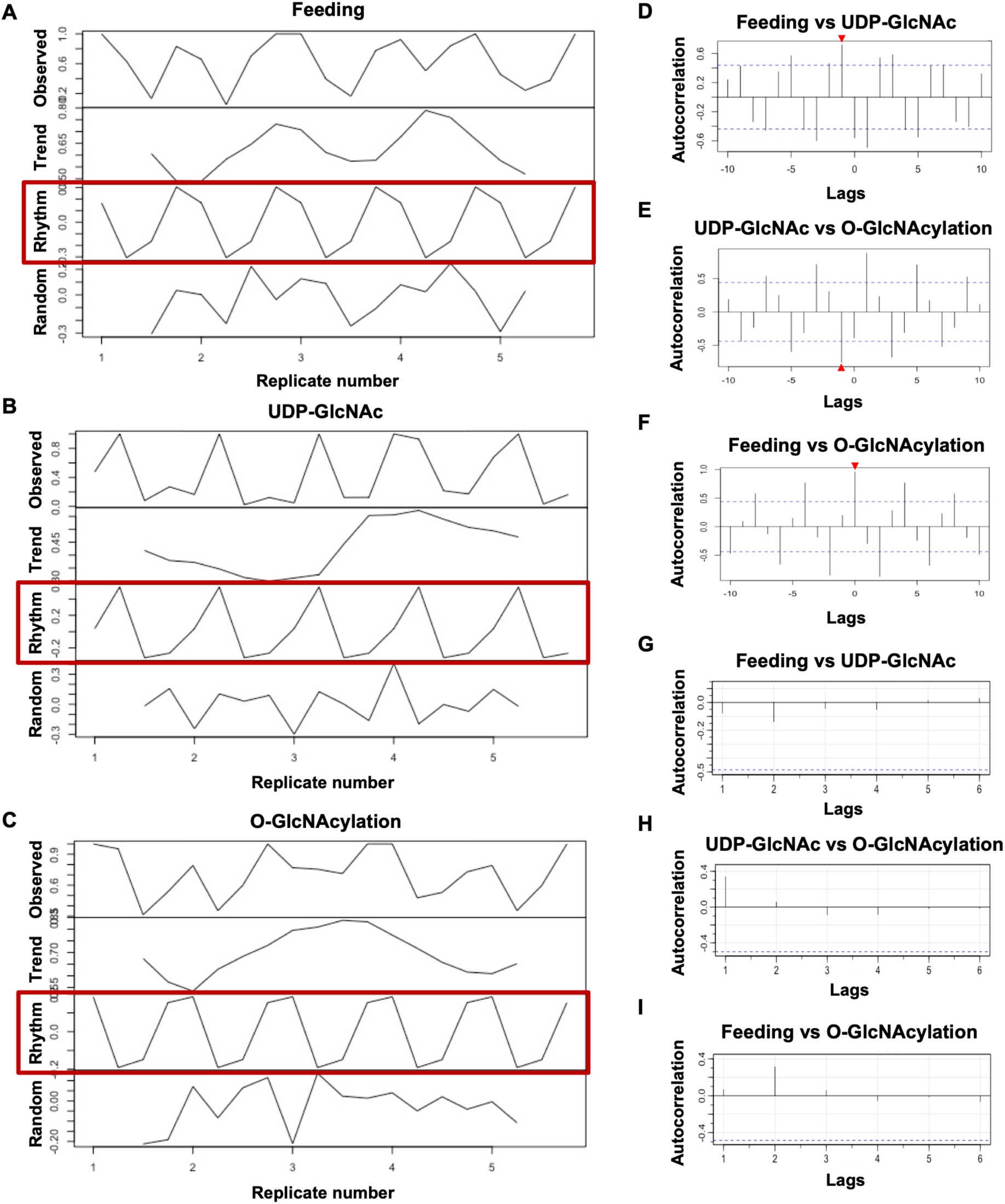
Cross correlation between the rhythms of feeding activity, UDP-GlcNAc, and nuclear protein O-GlcNAcylation. (A-C) Decomposition of observed raw data. Only rhythmic patterns of (A) feeding activity, (B) UDP-GlcNAc, and (C) O-GlcNAcylation were used for cross correlation analysis (boxed in red). (D-F) The calculation of cross correlation between rhythms. (D) UDP-GlcNAc rhythm is significantly correlated to feeding rhythm by lag -1 (i.e. The current UDP-GlcNAc level can be predicted from feeding activity that is around 6 hours earlier). (E) O-GlcNAcylation rhythm is significantly correlated to UDP-GlcNAc rhythm also by lag -1, while (F) O-GlcNAcylation rhythm is significantly correlated to feeding rhythm by lag 0. Dash lines indicate the cutoff of significance. Red triangle indicates the lag where two rhythms are significantly correlated with each other. (G-I) The correlation of residuals from the predicted models (Fig. 2F-H). All the residuals are not significantly correlated with each other, indicating our models successfully predicted all the possible correlation between rhythms. Dashed lines indicate the cutoff of correlation significance. Related to Figure 2.

**Figure S3.**
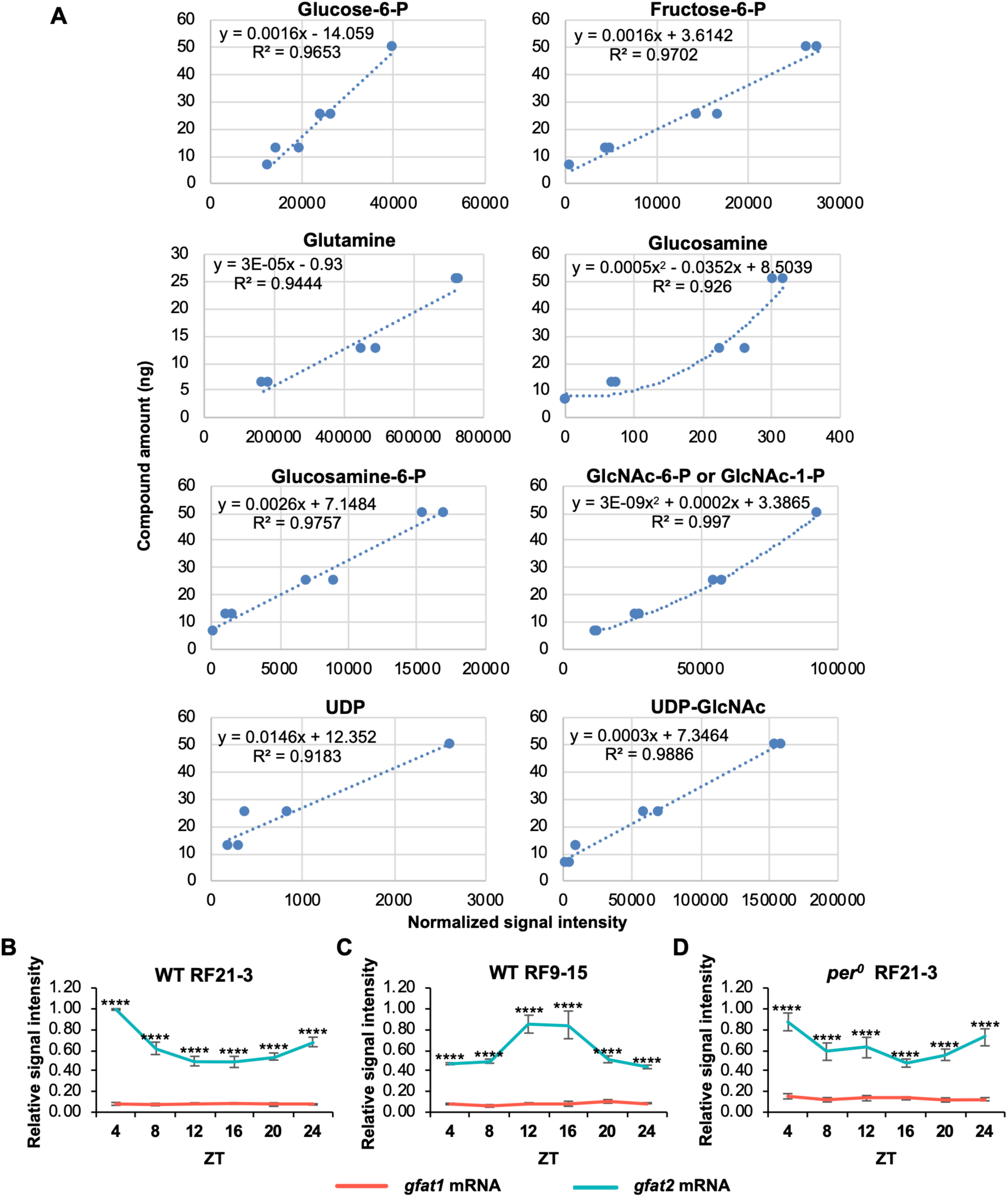
Analysis of HBP metabolites and *gfat* mRNA expression. (A) Standard curves of HBP metabolites generated for targeted metabolomics analysis. The peak intensity of HBP metabolites in HILIC-MS is proportional to their concentration. The formula for each of the trend curves is indicated on the top left corner of each panel. (B-D) The expression level of *gfat2* is 5- to 10-fold higher than that of *gfat1* in fly bodies. Line graphs to compare *gfat1* and *gfat2* mRNA levels in (B-C) WT (*w^1118^*) and (D) *per^0^* flies (n=3). Data are normalized (peak=1) and presented as mean ± SEM. Asterisks denote significant differences between *gfat1* and *gfat2* mRNA levels as determined by two-way ANOVA with *post-hoc* Tukey’s HSD tests. ****p<0.0001. Related to Figure 4.

**Figure S4.**
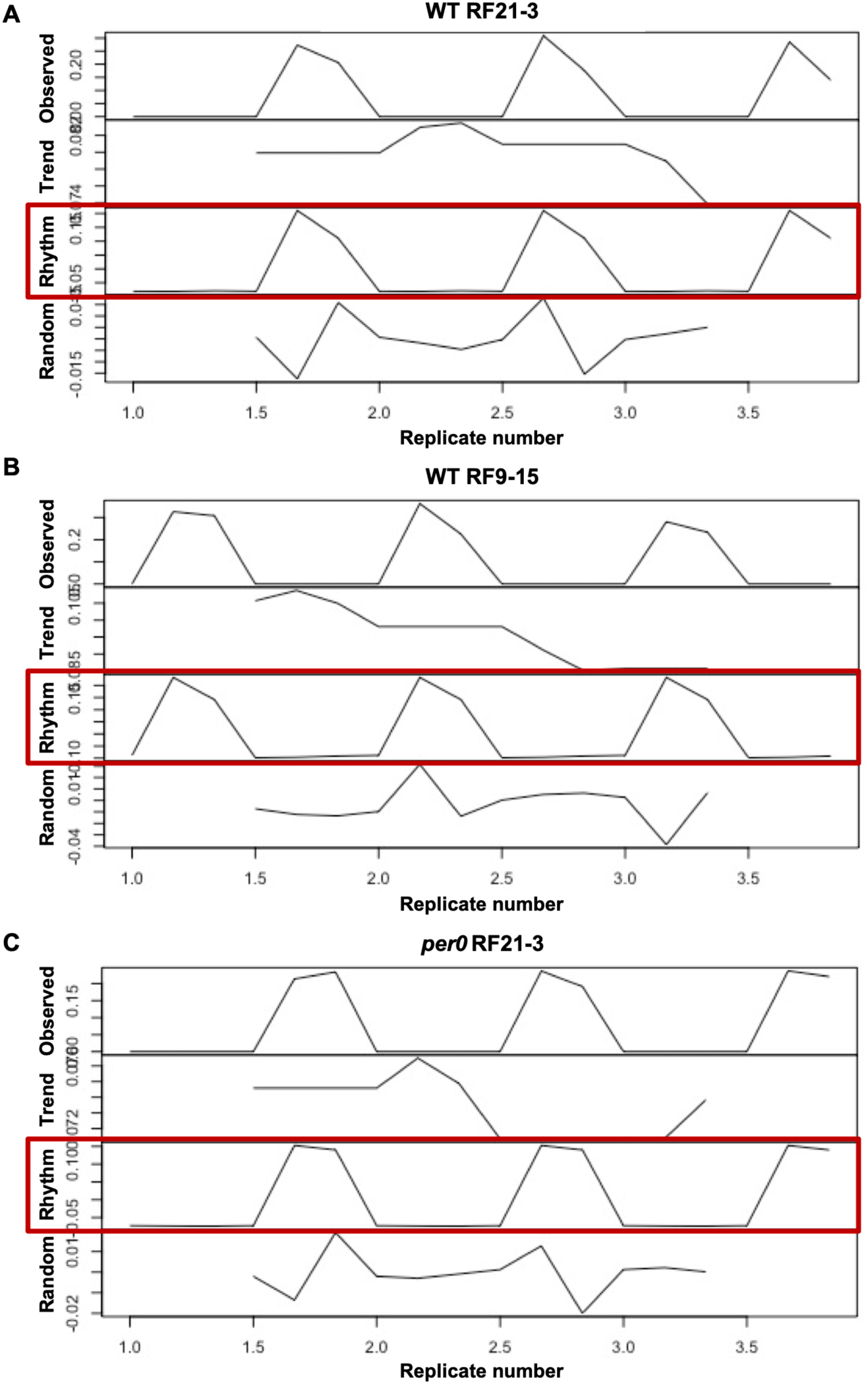
Decomposition of fly feeding data. The observed feeding data of (A) WT RF21-3, (B) WT RF9-15, and (C) *per^0^* RF21-3 groups were decomposed in R. Only rhythmic patterns of the feeding activity, denoted by red boxes, were used as input for the mathematical model. Related to Figure 6.

**Figure S5.**
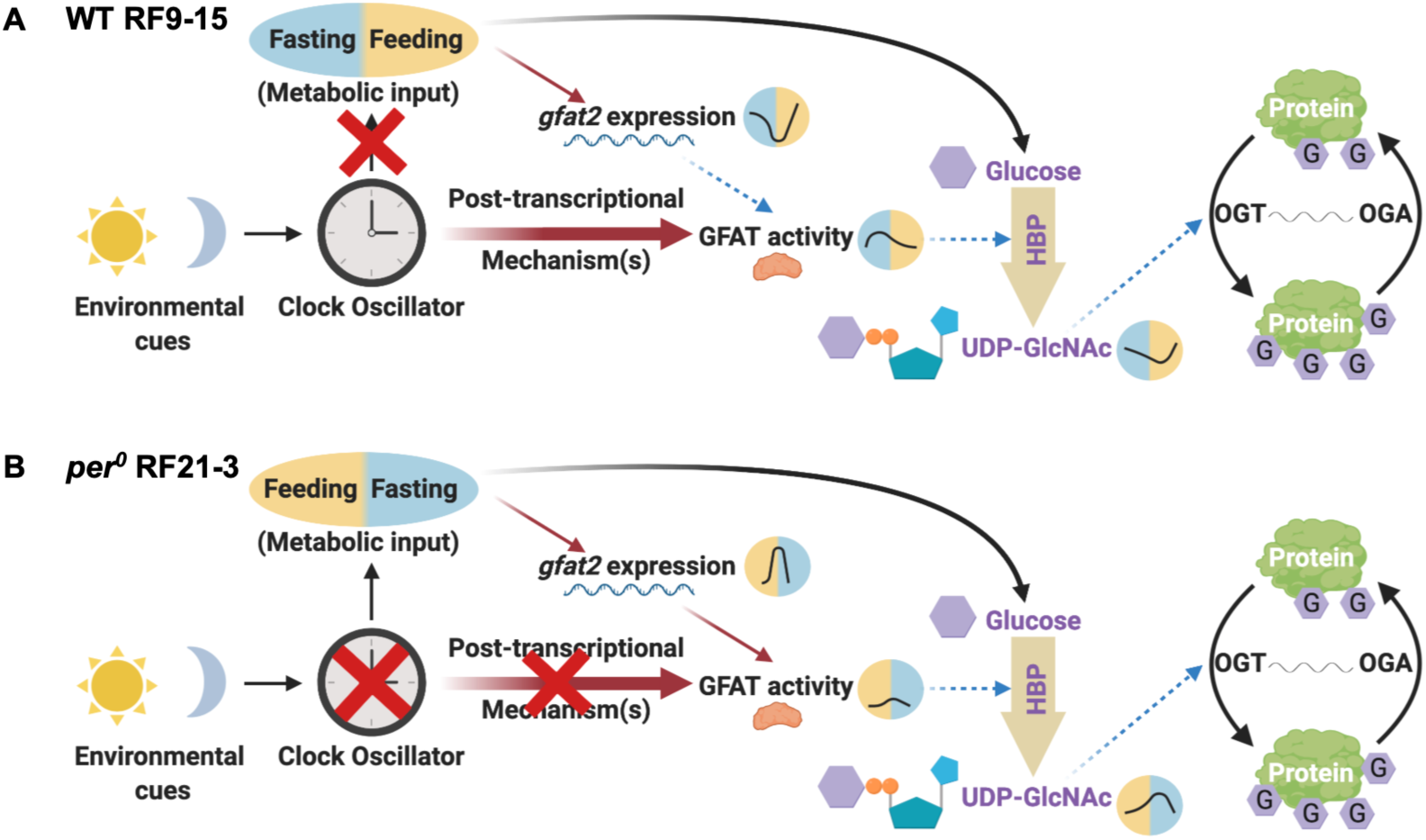
Schematic models showing the regulation of protein O-GlcNAcylation rhythm in WT RF9-15 and *per^0^* RF21-3 flies. (A) In WT RF9-15 flies, the post-transcriptional mechanism(s) still stimulates GFAT activity around their natural feeding time. However, the food-induced *gfat2* expression occurs later, which decreases its contribution to GFAT activity. Consequently, there is less nutrient flow through HBP and reduced production of UDP-GlcNAc, which causes the dampened protein O-GlcNAcylation rhythm. (B) In *per^0^* RF21-3 flies, clock-controlled post-transcriptional mechanism(s) is(are) absent. Although nutrient influx can still increase the expression of *gfat2* in a rhythmic manner, GFAT activity cannot reach a high level due to the lack of modulation by post-transcriptional mechanism(s). Consequently, the low GFAT activity after fly feeding results in lower UDP-GlcNAc production and dampened O-GlcNAcylation rhythm. Yellow background indicates feeding period, while the blue background indicates fasting period. Red arrows denote activation, and the thicker red arrows denote a stronger effect. The blue dashed arrows denote reduced effect of the indicated factors. Related to Figure 6.

**Figure S6.**
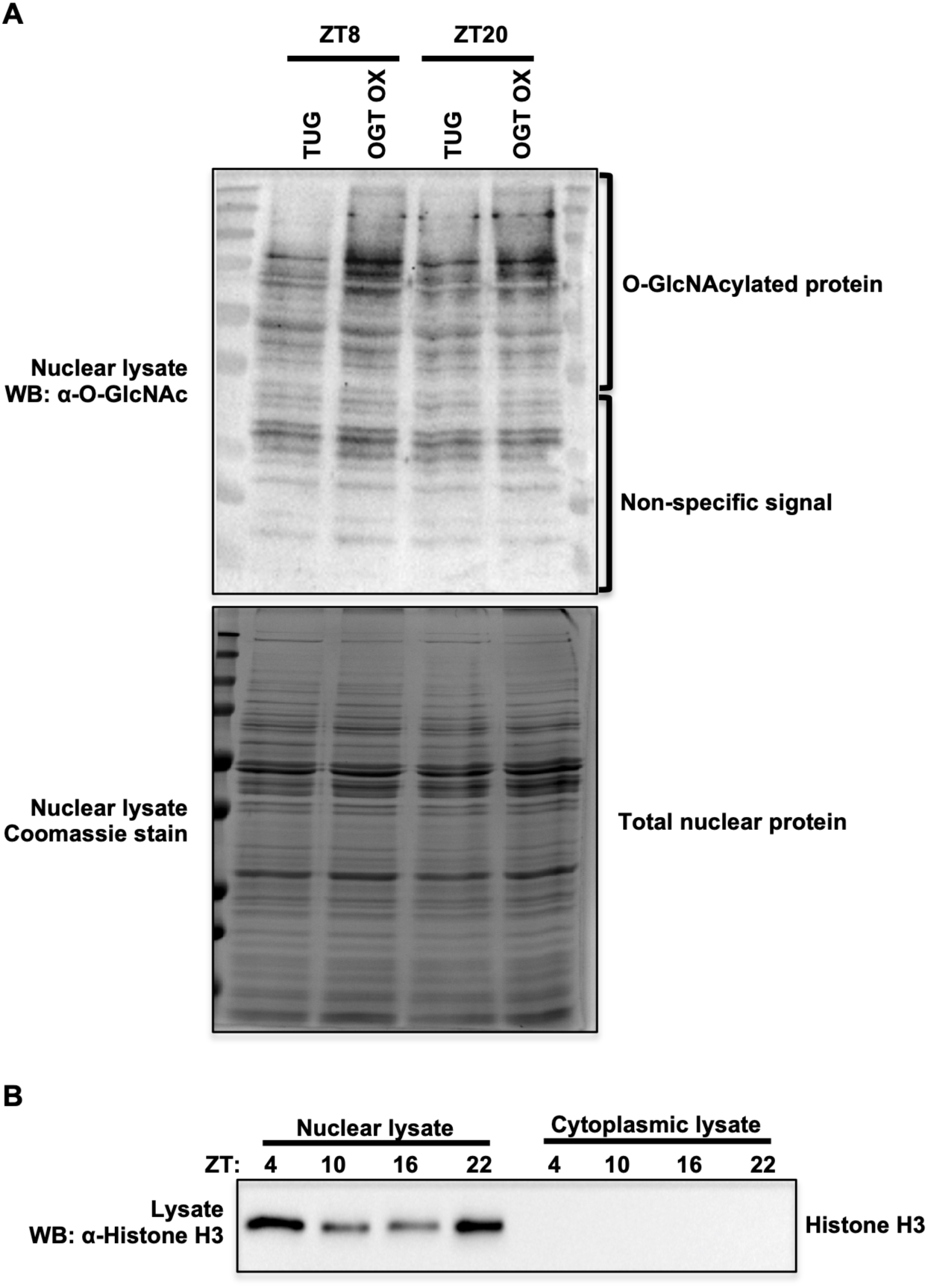
Validation of biochemical methods. (A) Validation of *α*-O-GlcNAc antibody to detect O-GlcNAcylated nuclear proteins in fly tissues. Flies overexpressing OGT in *timeless*-expressing clock neurons (*w; tim(UAS)-gal4; UAS-FLAG-ogt*) (Li et al., 2019), denoted as OGT OX, and parental control flies (*w; tim(UAS)-gal4*) were entrained in 12hrs light/ 12 hrs dark cycle at 25°C for 2 days and collected at ZT8 and 20 on LD3. Nuclear proteins were extracted from fly bodies and immunoblotting by *α*-O-GlcNAc was performed. Total nuclear proteins were stained by Coomassie blue (bottom panel) to indicate equal loading. (B) Validation of nuclear-cytoplasmic protein fractionation in wild type flies (*w^1118^*). Flies were entrained in 12hrs light/ 12 hrs dark cycle at 25°C for 2 days and collected at 4 time-points on LD3. Histone H3 was detected by immunoblotting to verify the purity of nuclear and cytoplasmic protein fractions. Related to Figures 1, 3, and 5.

**Table S1.** Circadian regulation of hexosamine biosynthetic pathway in mouse liver tissue based on published data.

| Enzymes | Transcripts | Protein | Phospho-peptide | Reference |
| --- | --- | --- | --- | --- |
| GFAT/GFPT | Cycling (2.3) | Non-cycling | Cycling (4.7) | [1-3] |
| GNPNAT | Cycling (9.4, 9.8) | Non-cycling | N/A | [2, 4-6] |
| PGM3 | Cycling (14.7) | Non-cycling | N/A | [2, 4, 7] |
| UAP1 | Cycling (2, 14.7) | Cycling (22) | N/A | [1-2, 4, 8] |

Abbreviations: GFAT/GFPT: Glutamine--fructose-6-phosphate aminotransferase; GNPNAT: Glucosamine-phosphate N-acetyltransferase; PGM3: Phosphoacetylglucosamine mutase; UAP1: UDP-N-Acetyl glucosamine pyrophosphorylase 1; N/A: Undetected. Numbers in parentheses denote peak phase of molecular rhythm.

**Table S2.**
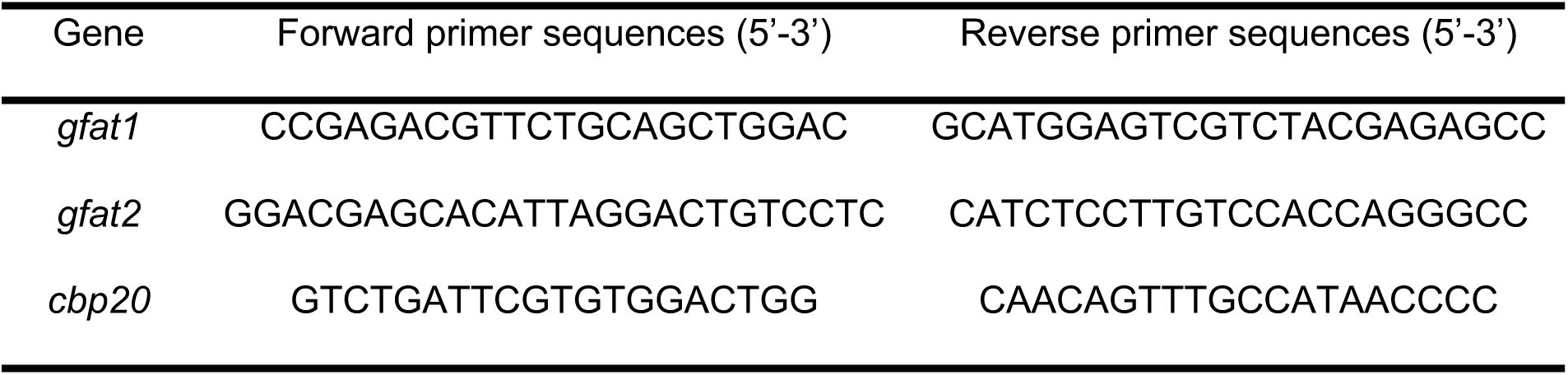
Primers for quantifying *gfat1* and *gfat2* mRNA

Dataset S1: Untargeted metabolomics of *Drosophila* heads and bodies on GC TOF platform

Dataset S2: Rhythmicity analysis (RAIN) of untargeted metabolomics

Dataset S3: Differential rhythm analysis (DODR) of HBP metabolites in TRF flies

